# Mitochondrial DNA variation of the striped hyena (Hyaena hyaena) in Algeria and further insights into the species’ evolutionary history

**DOI:** 10.1101/2025.03.13.643148

**Authors:** Louiza Derouiche, Mónica Rodrigues, Hafida Benameur-Hasnaoui, Ridah Hadj Aissa, Yasaman Hassan-Beigi, Seyed Massoud Madjdzadeh, Zuhair Amr, Aimee Cokayne, Paul Vercammen, Carlos Rodríguez Fernandes

## Abstract

The striped hyena (*Hyaena hyaena*) occurs in a wide range from north and east Africa, through southwest Asia to India, but its distribution is increasingly patchy and many of its populations are in decline due to intense human pressure. Its genetic diversity and structure, phylogeography, and evolutionary history, remain poorly understood. In this study, we investigated mitochondrial DNA variation in Algerian striped hyenas. Moreover, with the aim of contributing to our understanding of the evolutionary history of the species, we also examined samples from other geographic regions and compared our results with those of the only previous study in which individuals from across the range of the species were analysed. In particular, we performed a wide range of analyses of demographic history and estimation of the age of the extant mitochondrial DNA variation. The Algerian population sample was monomorphic. Overall, the global patterns of genetic diversity and the results of some demographic history analyses supported a scenario of population growth in the species, estimated to have occurred in the Late Pleistocene, but many of the analyses did not detect a significant signal of growth, most likely a result of the limited power provided by a small number of segregating sites. The estimates, from three different methods, for the time to the most recent common ancestor (TMRCA) of the mitochondrial DNA variation hovered around 400 ka, coinciding with one of the longest and warmest interglacials of the last 800,000 years, with environmental conditions similar to the Holocene.

## Introduction

The striped hyena *Hyaena hyaena* (Linnaeus, 1758), the only extant representative of its genus (Werdelin and Solounias 1991; Koepfli et al. 2006), is a large mammalian carnivore distributed from north and east Africa, through southwest Asia to India (Rieger 1981; Mills and Hofer 1998; Kingdon 2001; Holekamp and Kolowski 2009; Abi-Said and Dloniak 2015). Although once abundant and widespread across this range, its distribution is now mostly patchy, with the species extinct in many areas and the remaining populations generally in decline (Mills and Hofer 1998; Low Cunningham 2004; Kasparek et al. 2004; Khorozyan et al. 2011; Tourani et al. 2012; Abi-Said and Dloniak 2015; Akash et al. 2021; Bhandari et al. 2021; Çoğal et al. 2021). The global population is estimated at less than 10,000 individuals and, given the intense human pressure (persecution, poaching, poisoning, conflict with pastoralists, target of negative feelings and superstitious fears, prey base depletion, use as folk medicine, road mortality) to which it is subject, it is listed as ‘Near Threatened’ on the International Union for the Conservation of Nature (IUCN) Red List of Threatened Species (Abi-Said and Dloniak 2015).

The striped hyena is one of the four living species in the family Hyaenidae, the other three being the brown hyena (*Parahyaena brunnea*), the spotted hyena (*Crocuta crocuta*), and the aardwolf (*Proteles cristata*). The latter represents the oldest lineage of extant hyaenids, being quite distinct from the other three (Werdelin and Solounias 1991; Jenks and Werdelin 1998; Koepfli et al. 2006; Westbury et al. 2019). It has been estimated that the lineage ancestral to *Crocuta* diverged from the lineage ancestral to *Hyaena* and *Parahyaena* 9-11 Ma (Koepfli et al. 2006; Westbury et al. 2019, 2020), while the latter two lineages diverged from each other about 4-5 Ma (Rohland et al. 2005; Koepfli et al. 2006; Hu et al. 2021; Westbury et al. 2021). Traditionally, based on differences in body size and skull and pelage characteristics (Pocock 1934), the following five subspecies were accepted (Rieger 1979, 1981; Mills and Hofer 1998; Kasparek et al. 2004): *barbara* in Northwest Africa (type locality: Oran, Algeria); *dubbah* in East Africa (type locality: Atbara, Sudan); *syriaca* in Syria and Anatolia (type locality: Antakya, Turkey); *sultana* in southern Arabia (type locality: Mt. Qara, Oman); and *hyaena* from Iran to India (type locality: Benna Mountains, Larestan, Iran). However, the distributions and limits of these classically recognized subspecies are not precisely known (Rieger 1981), and attempts to delineate their geographic ranges and boundaries should be seen as tentative (Mills and Hofer 1998). Jenks and Werdelin (1998) noted that molecular data could be very useful for assessing subspecies designations in the striped hyena. In fact, a study that sequenced a 340-bp fragment of the mitochondrial Cytochrome b (*Cyt b*) gene in 13 striped hyena samples from scattered locations throughout the historical and current distribution of the species, found low genetic diversity (four haplotypes with two variable sites) and a general lack of obvious phylogeographic structure, with three of the four haplotypes shared between Africa and Eurasia (Rohland et al. 2005). Thus, although the species exhibits marked geographic variation in size and coat colour patterns, the currently accepted view is that there is no evidence for the recognition of subspecies (Kingdon 2001; Wozencraft 2005).

In Algeria, the striped hyena had the same history of decline over the past few centuries as observed in many other regions of its distribution, and the species remains vulnerable and threatened by anthropogenic causes of mortality and in need of conservation efforts, but it has managed to persist throughout much of the country (Kowalski and Rzebik-Kowalska 1991; Derouiche et al. 2020).

In this study, we sequenced a large sample of striped hyenas from Algeria (i.e., representing the traditional subspecies *barbara*) for a *Cyt b* fragment containing the shorter region analysed by Rohland et al. (2005) to allow comparisons. We also examined samples from geographic regions and traditional subspecies not included in that study, such as southern Arabia (subspecies *sultana*) and southern Iran (subspecies *hyaena*). Our objectives were to assess mitochondrial DNA (mtDNA) diversity of striped hyenas in Algeria and, expanding on the pioneering and seminal study of Rohland et al. (2005), contribute to increase our knowledge of the evolutionary history of the species.

## Materials and methods

### Sampling and laboratory procedures

To investigate the mtDNA diversity of the striped hyena in Algeria, we collected samples from 25 specimens found dead on roads or in the field. We also analysed samples of striped hyenas from Jordan (2), Saudi Arabia (3), Oman (5), and Iran (2) (Fig. 1 and Table 1). DNA was extracted from muscle tissue and blood samples using the EZNA Tissue DNA kit (Omega Bio-Tek), while that from dry hair samples was extracted following a simple polymerase chain reaction (PCR) buffer-based extraction protocol (Allen et al. 1998; Vigilant 1999). For every batch of samples, we used DNA extraction blanks to monitor contamination. We amplified a fragment containing the first 753 base pairs (bp) of the *Cyt b* gene with the primers L14724 (5’- TGATATGAAAAACCATCGTTG-3’; Irwin et al. 1991) and H15791 (5’- AATGTAGTTGTCTGGGTC-3’; Fernandes et al. 2008). For four hair samples (one from Algeria, one from Saudi Arabia, and the two from Jordan) we were unsuccessful in amplifying this fragment, which we interpreted as a result of DNA degradation. Thus, for those samples we amplified a 363-bp part of the fragment (corresponding to positions 16-378 of *Cyt b*), containing the 340-bp region analysed by Rohland et al. (2005), in two overlapping pieces, using respectively the two following primer pairs: Hyena Cytb F1 (5’- CCAATGACCAACATTCGA-3’)/Hyena Cytb R1 (5’-TCAGCCATAGTTGACGTC-3’) (for a fragment of 198 bp) and Hyena Cytb F2 (5’-AACCGCCTTTTCATCAGT- 3’)/Hyena Cytb R2 (5’-GACGTAACCTATGAATGC-3’) (for a fragment of 181 bp). PCRs were carried out in volumes of 15 μl with 1x PCR Buffer (NZYTech), 2 mM MgCl_2_, 0.2 mM of each dNTP (Bioline), 0.5 μM of each primer, 0.75 U of Supreme NZYTaq DNA Polymerase (NZYTech), and 3 μl of DNA extract. DNA extraction and PCR blanks were included to check for contamination. Thermal cycling conditions consisted of an initial denaturation at 95 °C for 5 min, followed by 45 cycles of 30 s at 94 °C, 30 s at 50 °C, 30 s at 72 °C, and a final extension of 7 min at 72 °C. The results of the PCR amplifications were visualized on 2% agarose gels to verify PCR quality and absence of contamination, and the PCR products were purified with an Exo-SAP protocol (Hanke and Wink 1994; Werle et al. 1994) and sequenced by the Sanger method at Macrogen Inc.

**Fig. 1.**
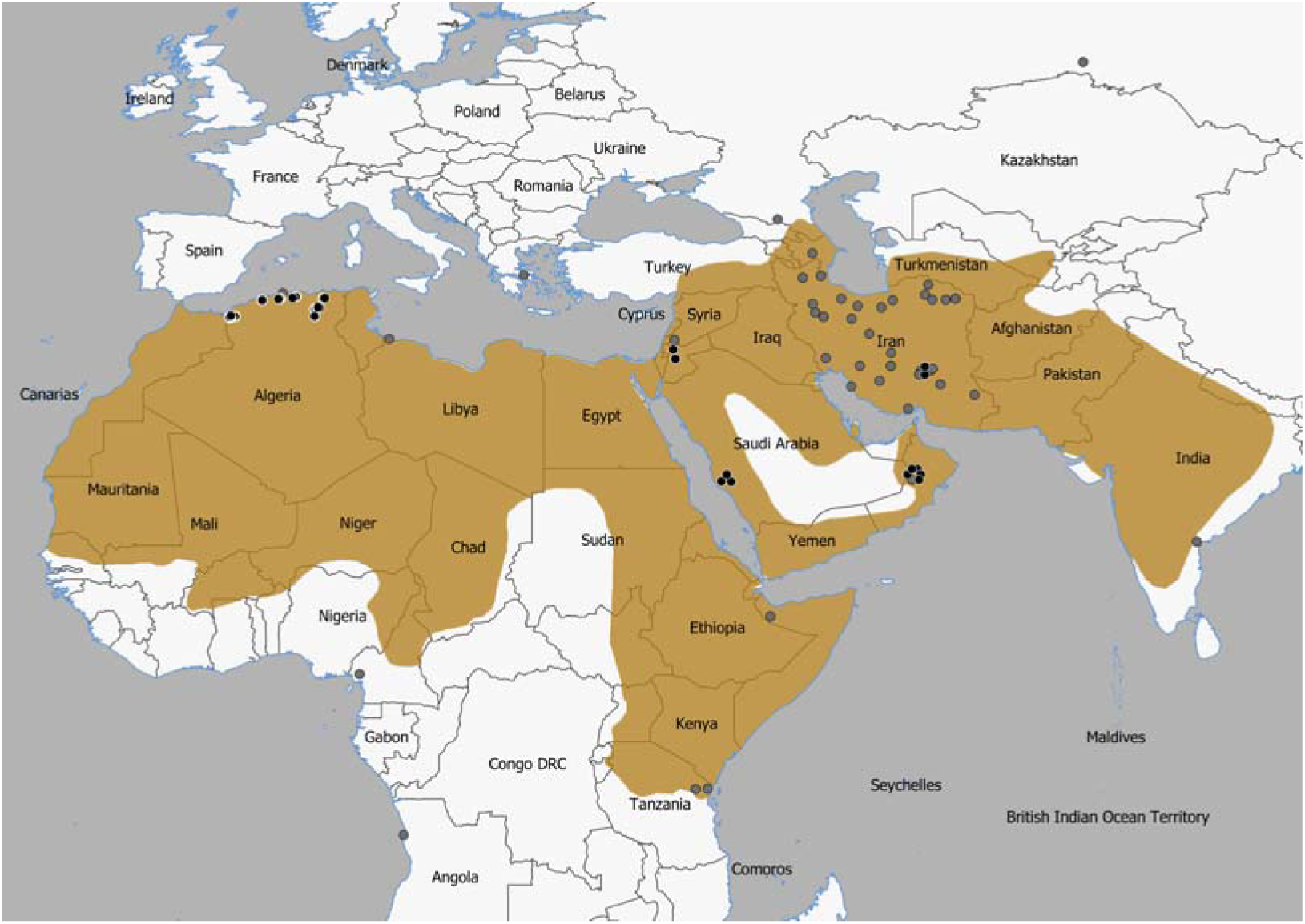
Map showing the geographic range of the striped hyena according to Abi-Said and Dloniak (2015). The plotted dots indicate the origin of the samples and sequences analysed; black dots in the case of our samples and dark grey dots for Genbank sequences (see Table 1 for details).

**Table 1.**
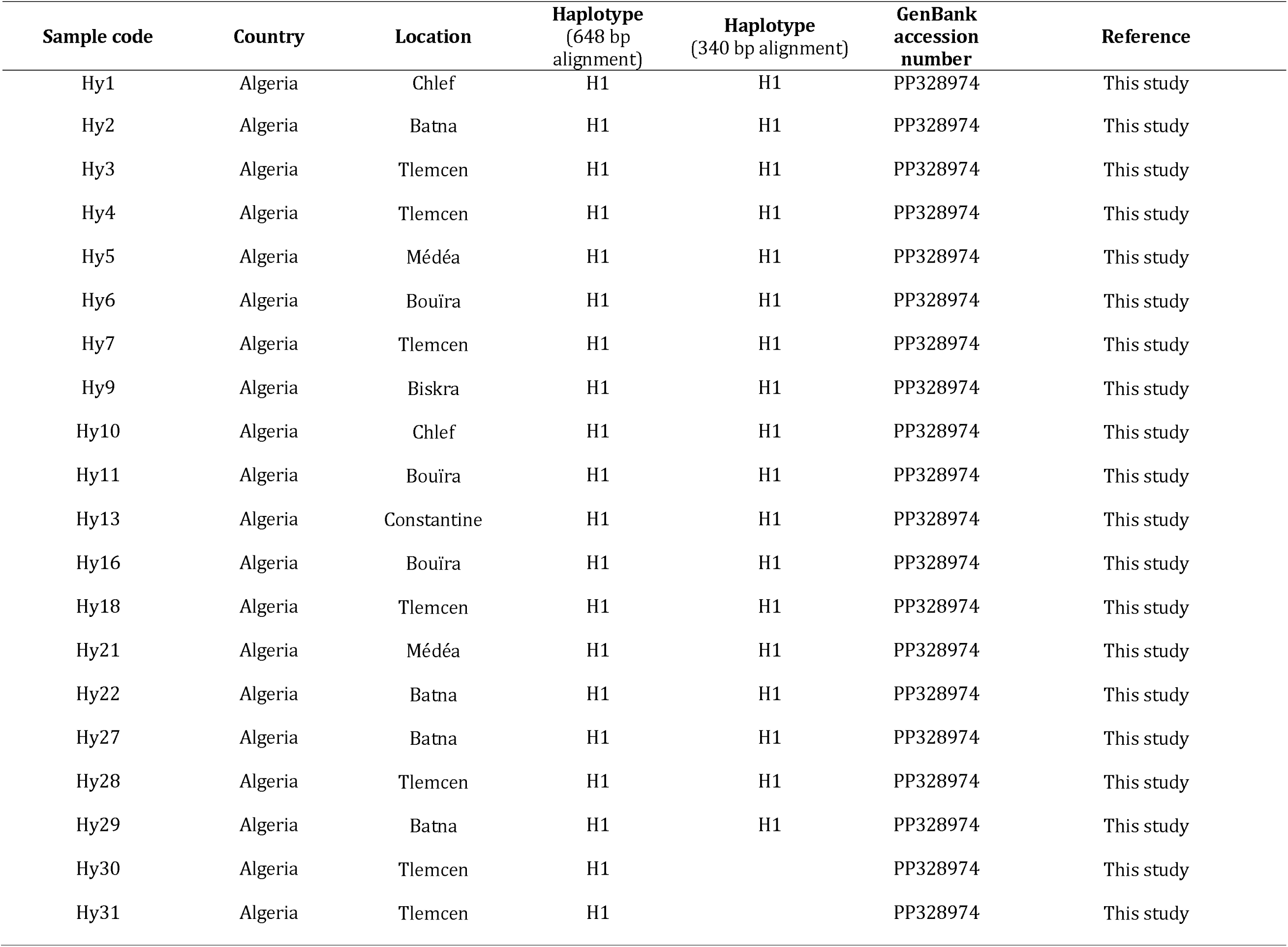
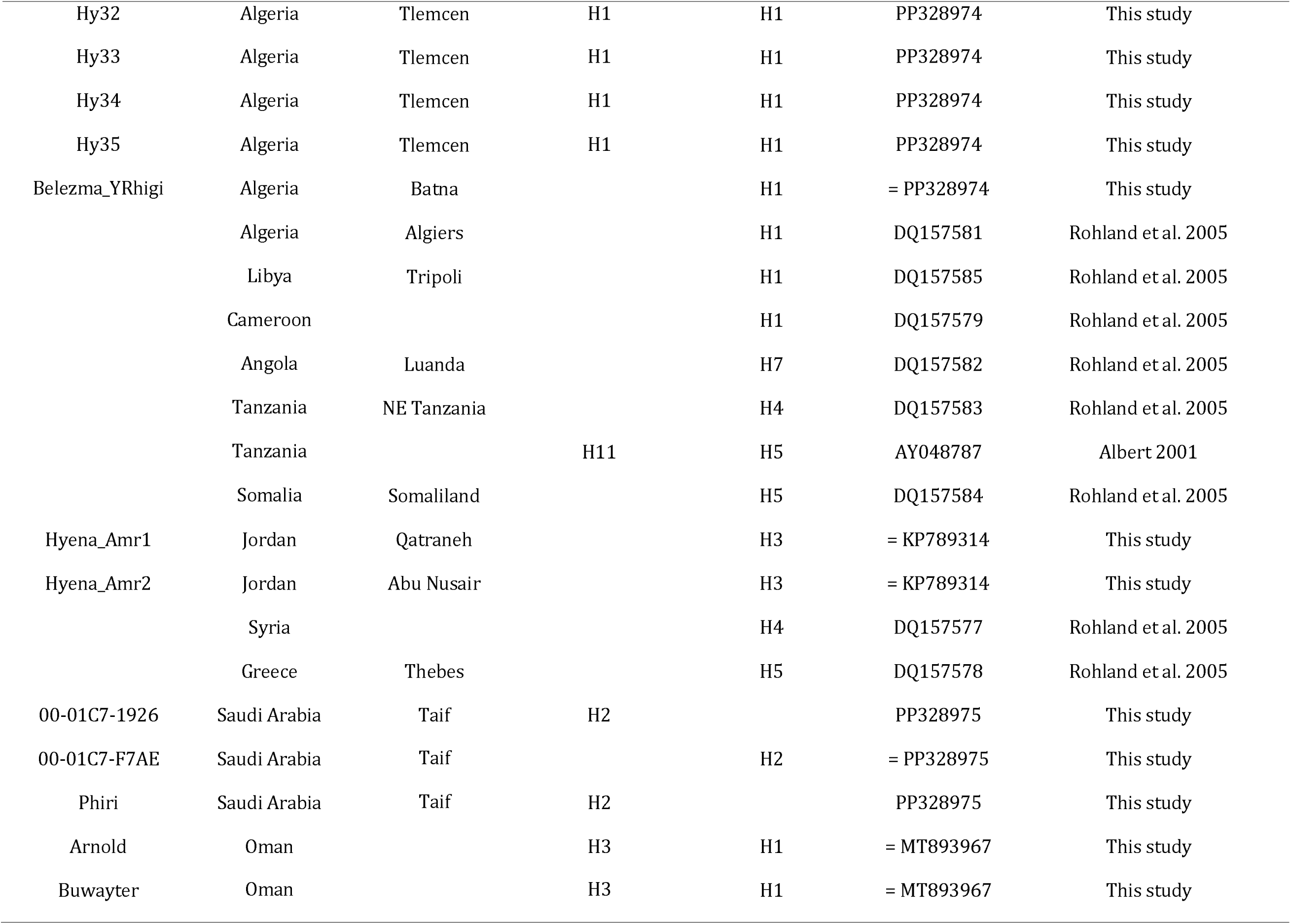
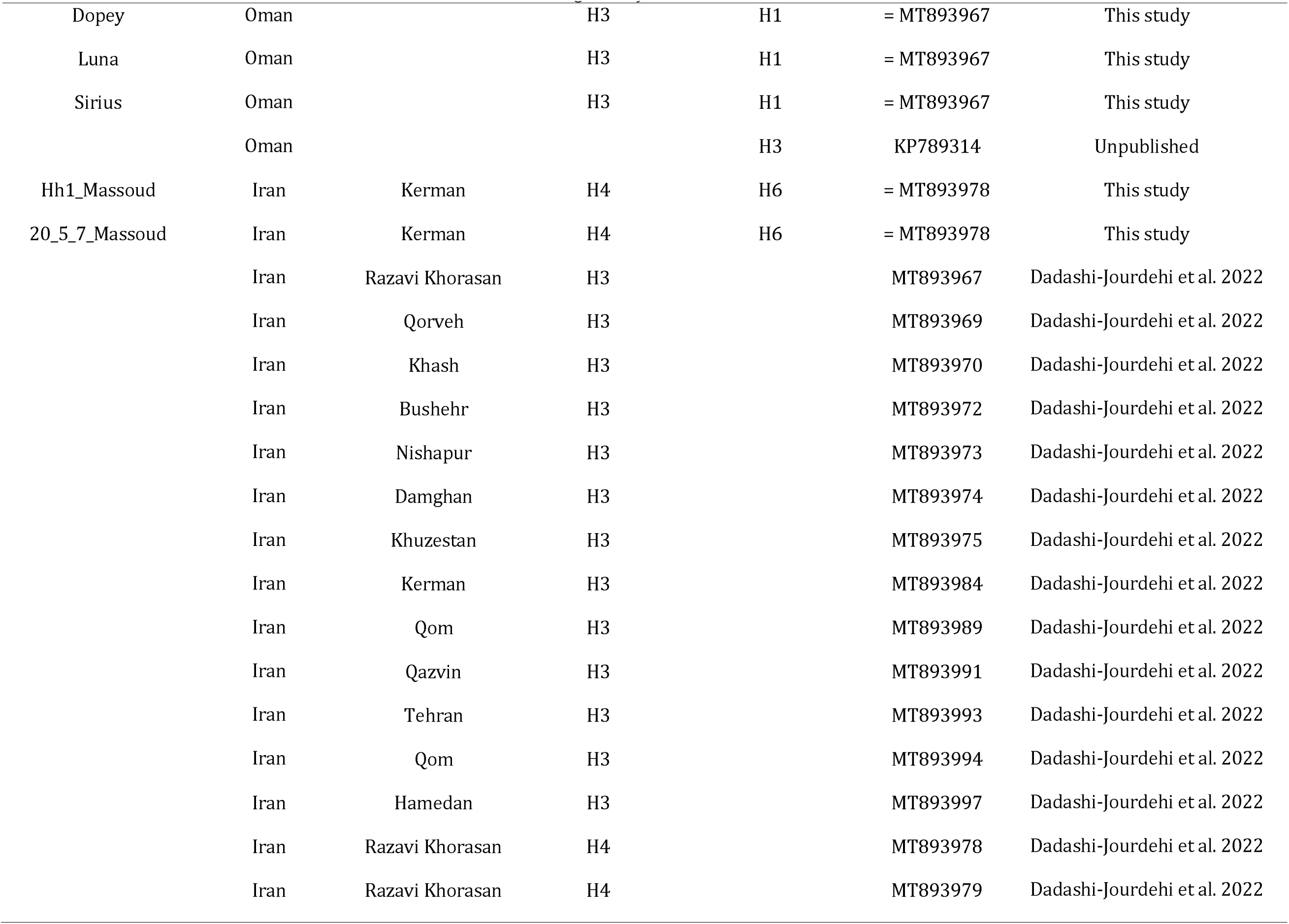
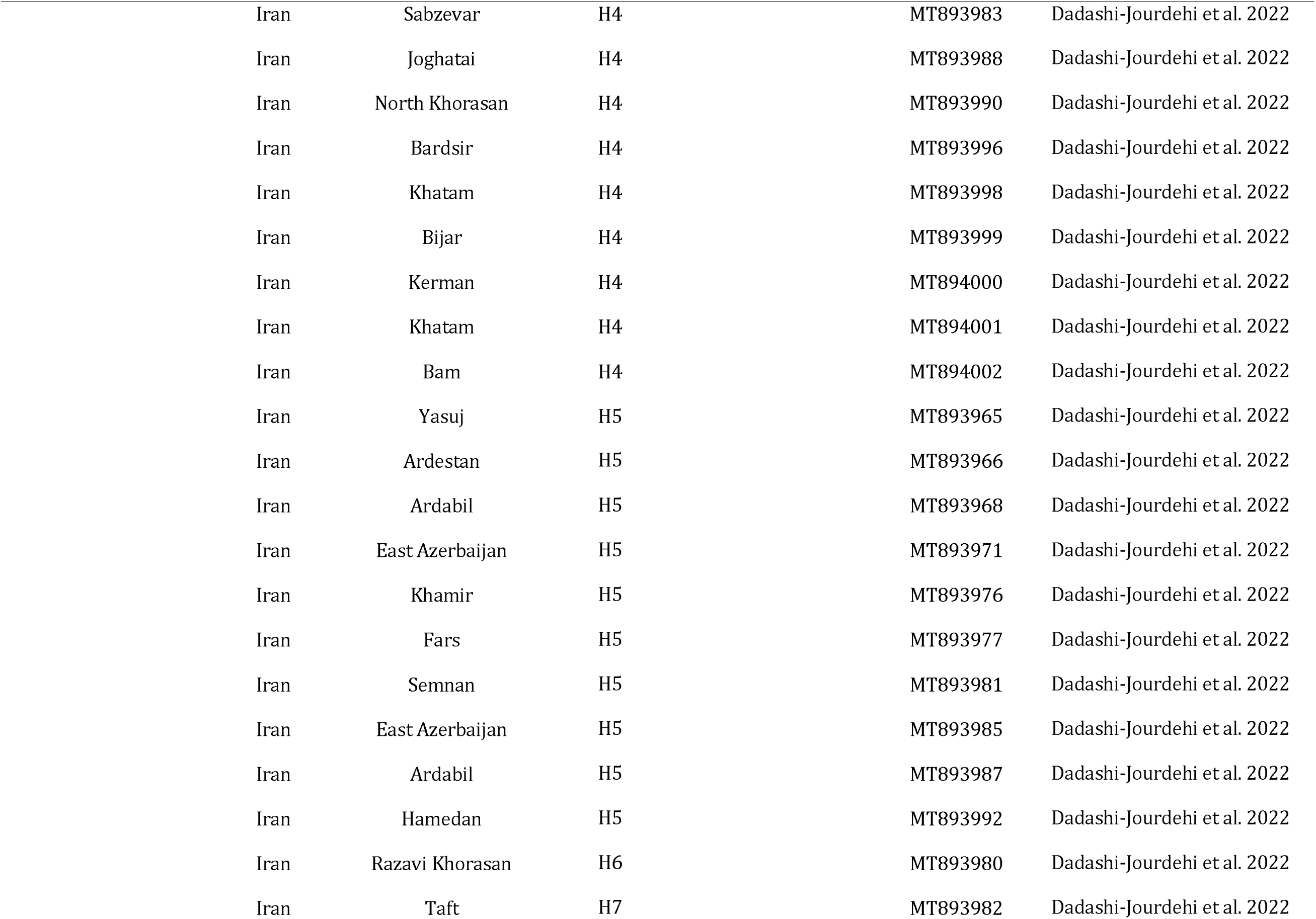
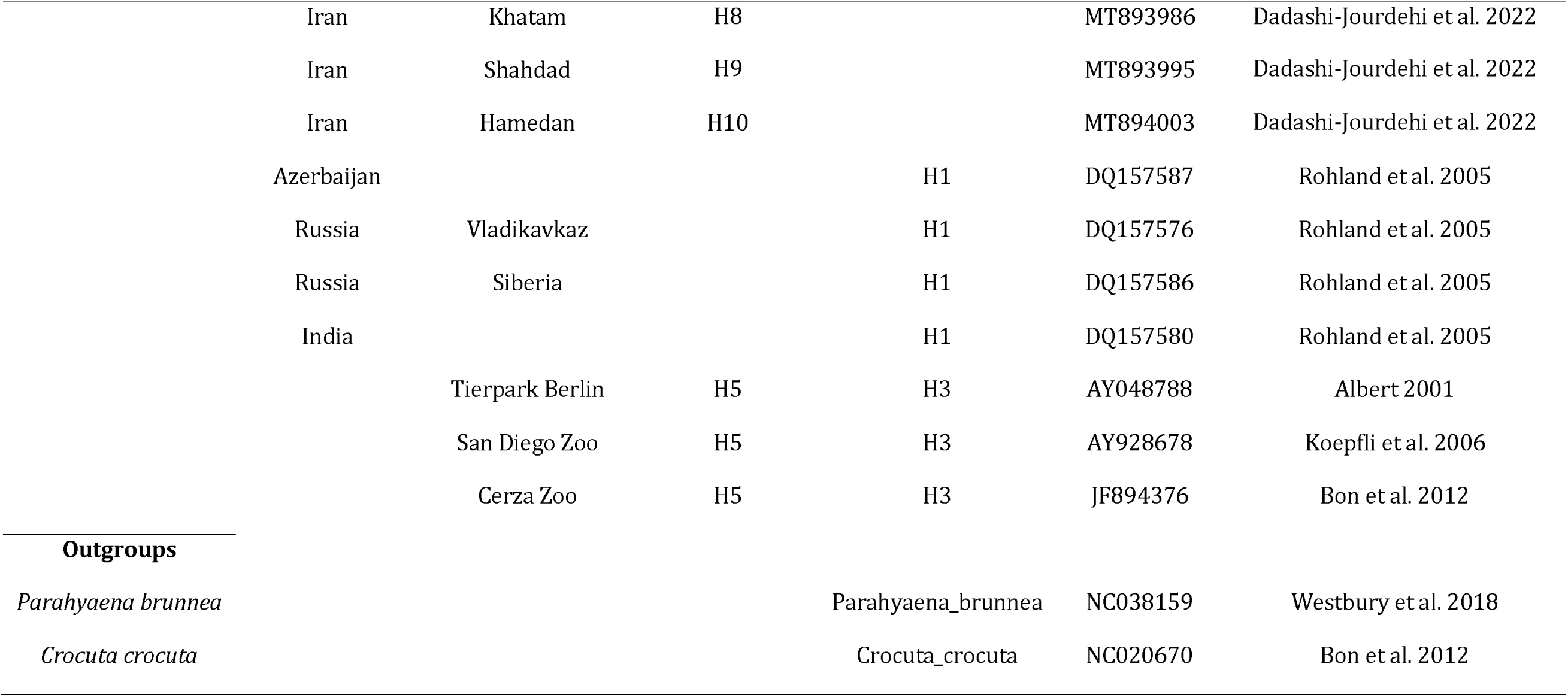
Information on the samples and previously published Cyt b sequences of striped hyenas and outgroups used in this study.

### Data analyses

Sequences were edited, assembled, aligned, and checked for the absence of indels and stop codons using SEQUENCHER 4.7 (Gene Codes Corporation) and the Translate program of the SEQUENCE MANIPULATION SUITE 2 (Stothard 2000). We downloaded striped hyena *Cyt b* sequences from GenBank (mainly the sequences from Rohland et al. (2005) and 39 sequences of Iranian individuals from Dadashi-Jourdehi et al. (2022)) and included these in our alignments (Table 1). Trimming our sequences to 648 bp allowed us to include those Iranian sequences from GenBank, but in order to incorporate other shorter sequences from diverse geographic origins, in particular those from Rohland et al. (2005), we created a smaller alignment of 340 bp (Table 1). Sequence alignments were analysed with FABOX 1.5 (Villesen 2007) to identify samples with identical sequences. File format conversions of sequence alignments for use in different computer programs were done using ALTER (Glez-Peña et al. 2010).

To characterize patterns of mitochondrial genetic diversity, and what they suggest in terms of demographic history of the striped hyena, we computed the number of polymorphic or segregating sites (S), the number of haplotypes (n_H_), haplotype diversity (h), nucleotide diversity (π), the mismatch distribution (Rogers and Harpending 1992; Schneider and Excoffier 1999), Tajima’s *D* (Tajima 1989), Fu’s *F_S_* (Fu 1997) and the *R_2_* statistic (Ramos-Onsins and Rozas 2002) in ARLEQUIN 3.5.2.2 (Excoffier and Lischer 2010) and DNASP 5.10.1 (Librado and Rozas 2009). We also computed Fu and Li’s (1993) D* and F* tests, as they are particularly powerful for detecting background selection (Fu 1997). If only these tests are significant, this suggests the action of background selection; if, on the contrary, they are not significant, but, for example, the Fu’s *F_S_* test is, then this is more compatible with a history of population growth (Fu 1997).

To visualize and examine the genealogical and geographical relationships among haplotypes, we estimated haplotype networks using the integer neighbour- joining method (IntNJ) in POPART 1.7 (Leigh and Bryant 2015). The IntNJ method was run with the reticulation tolerance parameter α set to zero. We used outgroup rooting (Jansen et al. 2002; Cassens et al. 2003; Dubach et al. 2013) to estimate the ancestral node of the ingroup. The outgroups used here, and in the phylogenetic tree reconstructions described next, were the brown hyena (*P. brunnea*) and the spotted hyena (*C. crocuta*).

To infer evolutionary relationships among haplotypes and assess the presence of clades within *H. hyaena*, we constructed phylogenetic trees. The tree reconstructions were conducted with the data partitioned by codon position, as this was the best-fit partitioning scheme according to the corrected Akaike information criterion (AICc) (Akaike 1974; Sugiura 1978; Hurvich and Tsai 1989) in PARTITIONFINDER 2.1.1 (Lanfear et al. 2017) using PHYML (Guindon et al. 2010). Phylogenetic analyses were performed using Bayesian inference (BI) and maximum likelihood (ML) as implemented in MRBAYES 3.2.6 (Ronquist et al. 2012) and IQ-TREE 1.6.12 (Nguyen et al. 2015), respectively. Analyses in MRBAYES were conducted with two parallel Markov Chain Monte Carlo (MCMC) runs, each with four Markov chains (one cold and three heated), default heating parameter (*t* = 0.1), and 20 million generations. The first five million generations were discarded as burn-in and, thereafter, chains were sampled every 500 generations. The entire general time-reversible (GTR; Lanave et al. 1984) substitution model space was sampled within the analyses (Huelsenbeck et al. 2004). Convergence was indicated by an average standard deviation of split frequencies between parallel runs of less than 0.01. For all model parameters, the effective sample size (ESS) was greater than 700 and the potential scale reduction factor (PSRF) was 1.0. Support for tree nodes was determined according to the values of Bayesian posterior probability (BPP) obtained in a majority-rule consensus tree (Holder et al. 2008; Huggins et al. 2011). In IQ- TREE we used the nucleotide substitution models estimated as best fit for each codon position, according to the Bayesian information criterion (BIC; Schwarz 1978), by ModelFinder (Kalyaanamoorthy et al. 2017), a model-selection method implemented in IQ-TREE (648 bp dataset, according to codon position: Kimura two-parameter model (K2P; Kimura 1980); Hasegawa-Kishino-Yano model (Hasegawa et al. 1985) with a proportion of invariable sites (HKY+I); HKY) (340 bp dataset, according to codon position: K2P+I; HKY; Tamura-Nei model (TN93; Tamura and Nei 1993)). Tree search runs were performed using IQ-TREE default settings, with the exception of the ’-allnni’ option which turns on a more thorough Nearest Neighbour Interchange (NNI) tree search, and support for each node was evaluated by 1000 non-parametric bootstrap replicates (Felsenstein 1985). Majority-rule consensus trees (Berry and Gascuel 1996; Holder et al. 2008) were computed with SUMTREES 4.5.2 of the DendroPy library version 4.5.2 (Sukumaran and Holder 2010) and visualized and edited with FIGTREE 1.4.4 (available at https://github.com/rambaut/figtree/releases).

To try to learn more about the recent evolutionary history of the species, we performed several analyses specifically on demographic history. In these analyses we used only the 648 bp dataset (76 striped hyenas; 13 segregating sites), as the 340 bp dataset (50 individuals) only contains five segregating sites. We constructed skyline plots (Ho and Shapiro 2011), which do not depend on a prespecified parametric model of demographic history, using the following four MCMC sampling-based methods: Bayesian Skyline Plot (BSP; Drummond et al. 2005), Extended Bayesian Skyline Plot (EBSP; Heled and Drummond 2008), Bayesian Skyride (Minin et al. 2008), and Bayesian Skygrid (Gill et al. 2013). These skyline plot methods were performed in BEAST 1.10.4 (‘Bayesian Evolutionary Analysis Sampling Trees’; Suchard et al. 2018), using the BEAGLE 3.1.0 library (Ayres et al. 2019), with the data partitioned by codon position. All BEAST input files were created in BEAUti, available in the BEAST package. Methodological details of the MCMC sampling-based skyline analyses are given in Supplementary Information. We compared different skyline models, and also simple parametric models (such as constant population size or exponential growth) (see Villanea et al. 2020), in terms of their relative fit to the data, using the Bayes factor (Jeffreys 1935; Kass and Raftery 1995), the ratio of the marginal likelihoods of the two models under comparison. The marginal likelihoods were estimated in BEAST with path sampling (Lartillot and Philippe 2006) and stepping-stone sampling (Xie et al. 2011), based on 100 path steps and a chain length of two million iterations. We also tested skyline plot methods based on a single tree (Ho and Shapiro 2011): classical (Pybus et al. 2000), generalized (Strimmer and Pybus 2001), and Bayesian multiple-change-point (MCP; Opgen- Rhein et al. 2005). The required ultrametric binary tree was a maximum clade credibility tree (MCCT) from an analysis in MRBAYES. MCMC in MRBAYES was performed as described above. The analysis used a strict clock model with a uniform prior probability distribution on branch lengths, a normally distributed clock rate prior with mean 0.01 and standard deviation 0.0025 (in substitutions per site per Ma) (how we obtained this substitution rate estimate is explained below), and a truncated normal prior probability distribution on the tree age with a mean of 0.4 Ma and a standard deviation of 0.15 (the choice of these values reflects the estimates obtained for the age of the mtDNA diversity, presented below). The single-tree skyline plot analyses were performed in the R package APE version 5.7.1 (Paradis and Schliep 2019). The analyses performed in the R environment used R version 4.2.3 (R Core Team 2023). Bayesian MCP analyses were performed assuming both a constant population size prior demographic function (Felsenstein 1992) and a skyline plot prior demographic function (Opgen-Rhein et al. 2005). We also estimated the fit of different demographic models (constant population size, exponential growth, expansion growth, logistic growth, with both continuous and piecewise variants) (Pybus and Rambaut 2002) to the MCCT in the R package genieR version 0.1.0 (Xiang et al. 2019).

We also used the Monte Carlo likelihood approach of Weiss and von Haeseler (1998), as implemented in IPHULA 1.16 (Schmidt et al. 2007), for inference on demographic history. In this analysis we used the HKY model, which was selected as best-fitting the 648 bp alignment by BIC in ModelFinder and also in JMODELTEST 2.1.10 (Darriba et al. 2012), and a transition/transversion rate ratio (kappa, κ) of 28 estimated in TREE-PUZZLE 5.3.rc16 (Schmidt et al. 2002). The method of Weiss and von Haeseler (1998) involves a model with three parameters (theta, θ: mutation-scaled effective population size (N_e_) in the past (‘initial’), i.e. before an eventual Ne change; tau, τ: mutation-scaled time since Ne started to change exponentially; rho, ρ: ratio of current to initial N_e_), which are estimated based on the number of segregating sites and the mean number of pairwise nucleotide differences (k). The method approximates the likelihood surface for the three parameters by coalescent simulations on a three-dimensional grid. After preliminary simulations to explore the likelihood surface and find the parameter ranges that enclose the global maximum likelihood peak, we ran 10 replicate runs, zoomed into grid areas of high likelihoods, for parameter estimation. In these final runs we defined five grid points, equally spaced between 1 and 20, for ρ, 10 grid points, equally spaced between 0.1 and 5, for θ, and 10 grid points, equally spaced between 0.1 and 2, for τ. We ran 50,000 simulations for each grid point (i.e. parameter combination), and the maximum allowed difference between simulated and observed k was set to less than 0.03 (i.e. ± 1% around the observed mean k of 1.72).

Inference on demographic history parameters was also performed with the MCMC coalescent genealogy sampler LAMARC 2.1.10 (Kuhner 2006; Kuhner and Smith 2007), assuming either an exponential growth or shrinkage model or a simpler model of constant population size. We used the Bayesian version of LAMARC, with a default starting value of θ per site of 0.01 and a logarithmic prior between 10-5 and 10; in the growth model analysis, we specified for the growth rate parameter (*g*) a starting value of 1 and a linear prior bounded between -500 and 5000. Percentile profile likelihoods were computed for both θ per site and *g*. The HKY model is not implemented in LAMARC, so we used the closest model available, Felsenstein84 (F84; Felsenstein and Churchill 1996), which in any case is very closely related to the HKY model. We ran four replicates of two final chains, in which for each chain we used an initial burn-in period of 200,000 discarded genealogies, followed by two million genealogies sampled every 50 for parameter estimation. We used adaptive heating, with four tree search temperatures (initial values: 1, 1.2, 1.5, 2) in each chain, and tree swapping between temperatures attempted at every step of the chain. TRACER 1.7.2 (Rambaut et al. 2018) was used to analyse the resulting trace files, assess run convergence, and calculate summary statistics and ESS values of parameters.

We also tested constant population size and exponential growth models in GENETREE 9.01, which implements a Markov chain technique for ancestral inference that simulates gene trees conditional on the mutation pattern in a sample of DNA sequences (Griffiths and Tavaré 1994a,b,c, 1995). GENETREE requires data compatible with the infinite sites model (Kimura 1969; Griffiths and Tavaré 1995). To meet this requirement, in the 648 bp dataset we needed to remove one polymorphic site (site 546) inconsistent with the model, which caused the haplotypes H1 and H3, as well as H7 and H10, to no longer differ from each other. In the constant population model, after preliminary runs with different values of the generating parameter for θ to optimize settings and identify the region of maximum likelihood of θ, we generated likelihood surfaces for θ based on 10 million simulations and 1000 points for the empirical distribution. In the exponential growth model, after preliminary runs to identify the likelihood surface regions of θ and growth rate yielding the highest joint likelihood, as the two parameters are not independent of each other, we generated likelihood surfaces separately for θ and growth rate, in both cases based on 100,000 simulations and 1000 points for the empirical distribution. A likelihood ratio test (LRT) was used to determine whether there was a significant difference in the fit to the data between the simpler constant population size model and the exponential growth model.

To estimate the time to the most recent common ancestor (TMRCA) of the observed mitochondrial genetic variation, we obtained an estimate of the substitution rate for the analysed *Cyt b* fragment as follows. Given our observed *Cyt b* divergence between striped hyena and its closest relative, the brown hyena, and the divergence between the two lineages that the fossil record suggests may have occurred 4-5.2 Ma ago (Werdelin and Solounias 1991; Werdelin et al. 1994; Werdelin 2003; Werdelin and Manthi 2012; Westbury et al. 2020, 2021), an estimate of the substitution rate can be obtained using equation *d_xy_* = 2μT (Nei 1985, 1987), where *d_xy_* is the mean sequence divergence per site, μ is the mean substitution rate per site, and T is the time since divergence. Genetic estimates of divergence time between *Hyaena* and *Parahyaena* also fall at about 4-5 Ma (Rohland et al. 2005; Koepfli et al. 2006; Hu et al. 2021; Westbury et al. 2021). We confirmed the assumption of nucleotide substitution rate homogeneity among the striped hyena haplotypes and between these and *Parahyaena*, using *Crocuta* as outgroup, via generalized relative rate tests with topological weighting (Robinson et al. 1998) in RRTREE 1.1.11 (Robinson-Rechavi and Huchon 2000), and with Tajima’s relative rate test (1993) as implemented in MEGA 11.0.9 (Tamura et al. 2021). In RRTREE, statistical significance (*P* < 0.05) of rate heterogeneity among lineages was assessed using both synonymous substitution rates (*Ks*) and nonsynonymous substitution rates (*Ka*). The *d*xy between striped hyenas and *Parahyaena* was computed in MEGA using the substitution model that best fits the data, according to the BIC in both IQ-TREE and MEGA, among the models available in MEGA for distance estimation (TN93 for the 648 bp alignment; K2P for the 340 bp alignment), and the standard error was estimated using 1000 bootstrap replicates. As a point value for the divergence time between *Hyaena* and *Parahyaena* we used 4.6 Ma, the midpoint in the 4-5.2 Ma interval suggested by the fossil record. Similar point values have been used or estimated for the divergence of the two lineages in recent genetic studies of hyaenids (Westbury et al. 2020; Hu et al. 2021; Westbury et al. 2021). The *d_xy_* estimate for the 648 bp alignment was 0.091 ± 0.013 (mean ± standard error), and for the 340 bp alignment was 0.086 ± 0.017 (mean ± standard error). Approximating 95% confidence intervals (CIs) based on 1.96 standard errors yields 95% CIs of [0.066, 0.116] and [0.053, 0.119], respectively. Using the above equation *d_xy_* = 2μT with T = 4.6 Ma, the *d_xy_* estimates yield substitution rate estimates of 0.99% per Ma (95% CI: 0.72-1.26%) and 0.93% per Ma (95% CI: 0.58-1.29%), respectively; i.e. ≈ 1% per Ma, the traditional average rate for mtDNA (Brown et al. 1979; Wilson et al. 1985). Also integrating the uncertainty in the fossil record about the divergence time between *Hyaena* and *Parahyaena* results, respectively, in the following CIs: 0.63-1.45% and 0.51-1.49%.

We estimated the TMRCA of the extant striped hyena mitochondrial genetic variation, based on both 648-bp and 340-bp datasets and using different methods implemented in Thomson’s estimator (Thomson et al. 2000), GENETREE and BEAST. An attractive feature of Thomson’s estimator is that it does not rely on a specific population genetic model (Thomson et al. 2000). On the other hand, the estimator requires data consistent with the infinite sites model. Accordingly, as mentioned above for the GENETREE analysis, in the 648 bp dataset we removed a polymorphic site (site 546). In the 340 bp dataset it was also necessary to remove one polymorphic site incompatible with the said model (site 317), which caused the haplotypes H1 and H7, as well as H4 and H5, to no longer differ from each other. We computed Thomson’s estimator and its variance (the latter according to equation 4 in Hudson 2007) using NumPy code kindly provided by Dr. Helmut Simon (https://github.com/helmutsimon), and 95% CIs (as in equation 5 in Hudson 2007). The required estimate of the unfolded site frequency spectrum was obtained using the site.spectrum function of the R package PEGAS (Paradis 2010) and an estimate of the ancestral sequence of the ingroup. We estimated this ancestral sequence using the two outgroups (*P*. *brunnea* and *C*. *crocuta*) in two different computer programs, FASTML (http://fastml.tau.ac.il/; Ashkenazy et al. 2012) and IQ-TREE, for comparison purposes, and both provided the same estimate of the ingroup’s ancestral sequence for each of the two datasets. In GENETREE, we used both the constant population model and the exponential growth model, with the ML estimates of θ and growth rate (obtained as described above) as generating parameters. Both the 648-bp and 340-bp datasets were made compatible with the infinite sites model by removing one inconsistent site in each dataset, as described above. The ingroup’s ancestral sequences used, one for each of the two datasets, were those mentioned above, estimated using FASTML and IQ-TREE. We ran 100 million simulations to obtain TMRCA estimates and likelihood surfaces with 10,000 points. In BEAST we ran analyses with both outgroups and only with *P. brunnea*. The outgroups correspond to the molecular clock calibrations used: i) the divergence between the *Crocuta* lineage and the lineage ancestral to *Hyaena* and *Parahyaena*, for which we used a normally distributed calibration prior with mean 9.5 Ma and standard deviation 2 Ma (Jenks and Werdelin 1998; Rohland et al. 2005; Westbury et al. 2019, 2020; Hu et al. 2021); and ii) the split between *Hyaena* and *Parahyaena*, with a normal calibration prior with mean 4.6 Ma and standard deviation 0.3 Ma (Rohland et al. 2005; Koepfli et al. 2006; Hu et al. 2021; Westbury et al. 2020, 2021). The datasets were partitioned by codon position. We used the piecewise-constant skyline tree prior (Drummond et al. 2005), which has been shown to perform well in molecular dating analyses involving a mixture of inter- and intraspecific data (Ritchie et al. 2017; Mello et al. 2021), and the approximate continuous time Markov chain (CTMC) rate reference prior (Ferreira and Suchard 2008). In the analyses of the 648 bp dataset with two outgroups, we used relaxed clock models (uncorrelated lognormal model, UCLN, Drummond et al. 2006; random local clocks model, RLC, Drummond and Suchard 2010) because a LRT (performed in MEGA) of the strict molecular clock hypothesis rejected it (P < 0.001). In contrast, this hypothesis was not rejected (P = 0.060) for the dataset with only *P. brunnea* as outgroup, in agreement with the results from RRTREE and Tajima’s relative rate test, and therefore we assumed a strict molecular clock in the analysis of this dataset. Similarly, the strict clock hypothesis was rejected for the 340 bp dataset with two outgroups (P = 0.007), but not for the one with only *P. brunnea* as outgroup (P = 0.226). For each BEAST analysis, we ran four independent replicate MCMC simulations, each with 50 million generations and sampled every 5000 generations following a pre-burnin of 10%. We also ran two replicate simulations without data to obtain estimates from the prior distribution and test the influence of priors on posterior distributions (Drummond et al. 2006; Brown and Smith 2018). For each analysis, we used TRACER with the default burn-in to assess convergence of the chain to the stationary distribution, obtain estimates and ESS of parameters, and plot marginal posterior densities. After verifying convergence, and confirming that posterior distributions of estimates were markedly different from the prior distributions, the tree files from the four runs with data were combined using SUMTREES, with the first 25% (2500) trees from each run discarded as burn-in, into a MCCT with median node ages. This method has a good overall performance, in terms of accuracy in estimating ages and model fit, compared to other tree summary approaches (Morrison 2008; Battistuzzi et al. 2010; Heled and Bouckaert 2013; Bromham et al. 2018). After analysing the results of the runs with the relaxed clock models, we noticed that in general the TMRCA estimates for the striped hyena mitochondrial variation seemed too high (> 800 ka) when compared with those obtained with Thomson’s estimator (range of mean estimates: ∼ 410-420 ka) and with GENETREE (range of mean estimates: ∼ 240-330 ka) and the only previously published estimate that we are aware of (340 ka; Rohland et al. 2005). Thus, we considered these high estimates likely to be erroneous. The apparent poor performance of the UCLN clock model does not seem to be due to the presence of significant rate autocorrelation among lineages, since, for both datasets with two outgroups, the 95% highest posterior density (HPD) interval of the covariance between parent and child branch rates contained zero (Drummond et al. 2006). The CorrTest method (Tao et al. 2019), implemented in MEGA, also did not reject the hypothesis of independent evolutionary rates among lineages (*P* > 0.05). However, we also noted that the standard deviation of the UCLN clock (ucld.stdev parameter) was low in the analyses of both datasets (648 bp dataset: mean = 0.049, median = 0.010, 95% HPD interval = [3.7 x 10-7, 0.243]; 340 bp dataset: mean = 0.138, median = 0.055, 95% HPD interval = [9.9 x 10-6, 0.557]), which suggests that the data is quite clock-like. The same indication was given by the results for the posterior distribution of the number of rate changes (*K*) (rateChangeCount parameter) in the RLC analyses, as a greater posterior than prior probability for *K* = 0 supports the global clock hypothesis, while small or negligible posterior probability for *K* = 0 strongly rejects the hypothesis (Drummond and Suchard 2010). In the analysis with the 648 bp dataset, the posterior distribution for *K* (mean: 0.619; median and mode: 0; 95 % HPD interval: [0,2]) showed a higher probability for *K* = 0 than the prior distribution. This was not so with the 340 bp dataset (mean: 0.769; median and mode: 1; 95 % HPD interval: [0,2]), for which 43% of the probability mass of the posterior distribution was for *K* = 1, but yet 42% of the probability was for *K* = 0. In such cases, a strict clock (SC) may perform better in dating estimation (Ho et al. 2005; Brown and Yang 2011). Marginal likelihood estimates, obtained using stepping-stone sampling in BEAST with 100 path steps and a chain length of 500,000, also supported the use of a SC (648 bp dataset: log marginal likelihood = -1344.5; 340 bp dataset: log marginal likelihood = -711.2), compared with the RLC (648 bp dataset: log marginal likelihood = -1344.5; 340 bp dataset: log marginal likelihood = -711.1) and the UCLN (648 bp dataset: log marginal likelihood = - 1347.4; 340 bp dataset: log marginal likelihood = -713.2). Therefore, we decided to also use the SC in the analyses with two outgroups.

## Results

### Genetic variation, haplotype relationships and phylogenetic analyses

The new striped hyena DNA sequences discovered in this study were deposited in GenBank under accession numbers PP328974-PP328975 (Table 1). Several observations indicate that the generated sequences are mitochondrial and not nuclear-integrated copies of mtDNA (Arctander 1995; Zhang and Hewitt 1996; Bensasson et al. 2001). First, PCRs consistently yielded single products of the expected size. Second, independent replicate PCRs produced identical sequences.

Third, the sequences were unambiguous and did not contain indels or stop codons (Smith et al. 1992; Mirol et al. 2000; Triant and DeWoody 2007).

Genetic diversity measures and results of mutation-drift equilibrium tests, for both the 648 bp and 340 bp alignments of striped hyenas, are presented in Table 2. In the longer alignment the number of polymorphic sites was 13, with nine being parsimony-informative and four singletons, and in the shorter alignment the number of polymorphic sites was five, four of them parsimony-informative and one a singleton. In both datasets, none of the mutation-drift equilibrium tests used rejected at the 95% confidence level the null hypothesis of a constant size population under the neutral model; note that in the *F*S statistic test, an α = 0.02 corresponds to a 5% significance level (Fu 1997). The D* and F* statistics were respectively -0.739 (P = 0.233) and -0.983 (P = 0.180) for the 648 bp alignment, and 0.134 (P = 0.529) and -0.228 (P = 0.396) for the 340 bp alignment. Analysis of the mismatch distribution for the 648 bp dataset under a model of sudden demographic expansion (Schneider and Excoffier 1999), implemented in ARLEQUIN, did not reject the expansion hypothesis (P = 0.678; 20,000 bootstrap replicates). In agreement, the Harpending’s (1994) raggedness index was low (0.032; P = 0.552). The model parameter for the expansion time was estimated at 1.877 (in mutational time units) (95% CI: 0.715-3.736), which, assuming a nucleotide substitution rate of 1% per Ma, translates into 144,830 years (95% CI: 55,170-288,272). For the 340 bp dataset, the analysis of the mismatch distribution under the sudden expansion model failed to converge, possibly due to the low number of polymorphic sites in this dataset. However, analyses in DNASP showed that the observed mismatch distribution fits better with the expected distribution under a sudden expansion model (Rogers and Harpending 1992) than under a model of constant population size (Fig. S1). We also attempted in ARLEQUIN to test the mismatch distribution under a model of spatial expansion (Ray et al. 2003; Excoffier 2004), but the least-square procedure to fit the expected and observed mismatch distributions did not converge.

**Table 2.** Estimates of mitochondrial genetic diversity and mutation-drift equilibrium tests in striped hyenas, based on the two Cyt b alignments with different sequences and lengths n, number of samples; S, number of segregating sites; nH, number of haplotypes; h, haplotype diversity (mean and standard deviation); π, nucleotide diversity (mean and standard deviation); D, Tajima’s D statistic (Tajima 1989); F_S_, Fu’s F_S_ statistic (Fu 1997); R_2_, Ramos-Onsins and Rozas’ R_2_ statistic (Ramos-Onsins and Rozas 2002); P, p-value of the test.

|  | **S** | **n<sub>H</sub>** | **h** | **π** | **D** | **F<sub>s</sub>** | **R<sub>2</sub>** |
| --- | --- | --- | --- | --- | --- | --- | --- |
| 648 bp dataset (n = 76) | 13 | 11 | 0.794 ± 0.022 | 0.0026 ± 0.0017 | -0.993<br>(P = 0.166) | -2.662<br>(P = 0.126) | 0.064<br>(P = 0.181) |
| 340 bp dataset (n = 50) | 5 | 7 | 0.498 ± 0.082 | 0.0020 ± 0.0017 | -0.922<br>(P = 0.198) | -3.345<br>(P = 0.020) | 0.071<br>(P = 0.149) |

Both the haplotype networks (Fig. 2) and the BI and ML phylogenetic trees (Figs. S2-S4), for both the 648 bp and 340 bp datasets, indicate a shallow genetic divergence among the striped hyena haplotypes, and essentially an overall lack of phylogeographic structure.

**Fig. 2.**
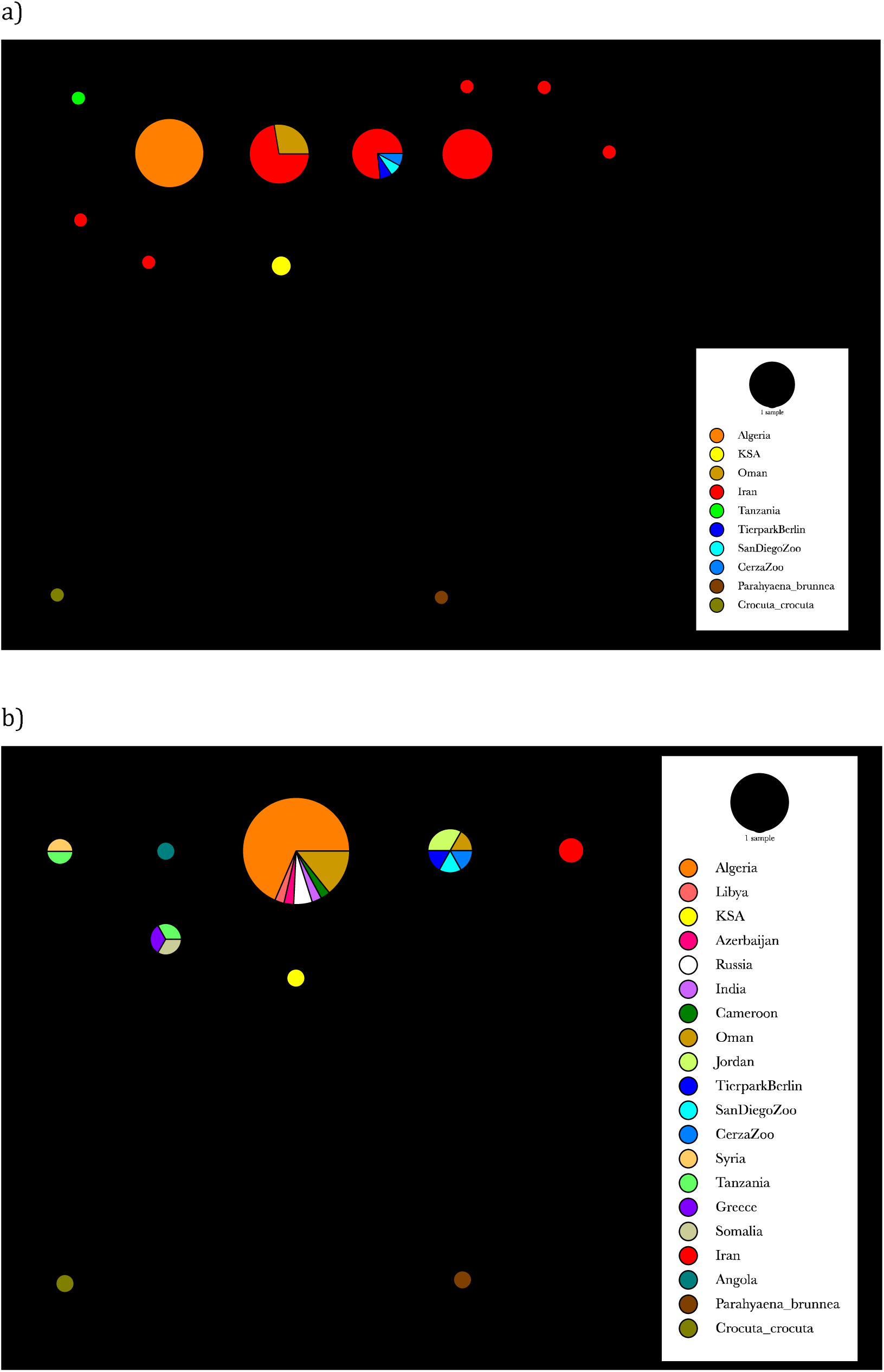
IntNJ networks of (a) 648-bp and (b) 340-bp striped hyena Cyt b haplotypes, rooted with two outgroups (brown hyena and spotted hyena). The first contains 76 striped hyena sequences and the second contains 50. Circles represent haplotypes and their size is proportional to frequency. Circles are coloured according to where haplotypes were found and to their relative frequency. Small black circles represent hypothetical haplotypes. Dashes on lines connecting haplotypes represent the number of nucleotide substitutions separating them. Haplotype designations and further information are given in Table 1.

### Demographic simulations

The different MCMC sampling-based skyline plot analyses conducted in BEAST did not identify signs of significant population size change in the Late Quaternary demographic history of the striped hyena. In the BSP analyses using a normally distributed prior for the substitution rate with mean 0.01 and standard deviation 0.0025 (in substitutions per site per Ma), truncated to be positive (corresponding to a 95% interquantile range of 0.005-0.015), the estimated mean tree root height was 281 ka (median: 248 ka; 95% HPD interval: 61-571 ka) for the piecewise-constant model (ESS > 1500 for all parameters), and 264 ka (median: 232 ka; 95% HPD interval: 60-536 ka) for the piecewise-linear model (ESS > 2200 for all parameters). The respective BSPs for the two models are shown in Fig. S5, in which it can be observed that the rise of the population size curve at the left end of the plots never exceeds the 95% HPD limits along the rest of the graph (Ho and Shapiro 2011; Grant 2015), where the curve is essentially flat. The results of analyses using either a uniformly or exponentially distributed prior for the substitution rate exhibited a similar pattern of no significant population size changes (not shown). In the EBSP analyses using the same normally distributed prior for the substitution rate as above, the estimated mean tree root height was 363 ka (median: 331 ka; 95% HPD interval: 85-717 ka) for the stepwise model (ESS > 2200 for all parameters), and 341 ka (median: 304 ka; 95% HPD interval: 58-688 ka) for the linear model (ESS > 2800 for all parameters). For the ’population size changes’ parameter in each model, the estimated mean was 0.867 (median and mode were 1, and the 95% HPD interval was 0-2) and 1.024 (median and mode were 1, and the 95% HPD interval was 0- 3), respectively. The respective EBSPs for the two models are presented in Fig. S6. Results of analyses using a uniformly distributed prior for the substitution rate also did not indicate significant population size changes in recent demographic history (not shown). The skyride (ESS > 6500 for all parameters) and skygrid (ESS > 650 for all parameters) analyses also yielded the same result (Fig. S7); their respective mean tree root height estimates were 72 ka (median: 66 ka; 95% HPD interval: 24-133 ka) and 277 ka (median: 245 ka; 95% HPD interval: 67-564 ka). In the analyses with the constant population size (uniform population size prior; ESS > 4700 for all parameters) and exponential growth (uniform population size and growth rate priors; ESS > 1500 for all parameters) models, the estimated mean tree root heights were 419 ka (median: 349 ka; 95% HPD interval: 111-809 ka) and 508 ka (median: 349 ka; 95% HPD interval: 103- 814 ka). The 95% HPD interval of the posterior distribution of the exponential growth rate parameter ([-1.803, 4.018]) included zero, i.e. did not reject a constant population size. The skygrid model was the one with the highest log marginal likelihood estimate (stepping-stone sampling MLE = -1096; path sampling MLE = -1095), but not significantly different from that of the piecewise- linear version of the BSP (MLE = -1097 by both stepping-stone sampling and path sampling). Still using Bayes factors, the skygrid model had positive support against the piecewise-constant version of the BSP (stepping-stone sampling MLE = -1099; path sampling MLE = -1098) and very strong support (2 ln Bayes factor > 10) against other models. The results of the single-tree skyline plot analyses done in APE suggested population growth during the Late Pleistocene (Figs. S8-S9). The fit of the different demographic models tested in genieR was compared using the Akaike information criterion (AIC). The constant population size model was rejected (ΔAIC = 90), while the exponential growth (AIC = -1575.919) and expansion growth (AIC = -1573.919) models were equally supported (ΔAIC ≤ 2).

The implementation of Weiss and Von Haeseler’s (1998) method in IPHULA estimated a maximum likelihood model parameter combination (ρ = 10, θ = 2.8, τ = 0.3) that suggests slow exponential growth since the Late Pleistocene. Specifically, assuming a nucleotide substitution rate of 1% per Ma and a generation time for the striped hyena of 9.2 years (Pacifici et al. 2013), a τ of 0.3 translates to about 23 ka. In turn, a θ of 2.8 corresponds to a female Ne of 23484 at that time. However, the 95% confidence set of models, based on the likelihood ratio of each combination of demographic parameters tested and the maximum likelihood model, included scenarios of constant population size.

In LAMARC, analyses using the exponential growth or shrinkage model suggested a slow exponential growth scenario. The highest likelihood estimate of *g* was 910 (95% HPD interval: [-83, 4704]; ESS > 700 in all individual replicate runs). Therefore, the 95% HPD interval crossed zero, not ruling out scenarios of constant population size or even contraction. Moreover, LAMARC estimates of *g* tend to be biased upwards when only a few loci are analysed (Kuhner et al. 1998). The highest likelihood estimate of current θ per site was 0.008 (95% HPD interval: [0.002, 0.020]; ESS > 350 in all individual replicate runs), which corresponds to a θ of 5.2 for the 648 bp fragment and, again assuming a nucleotide substitution rate of 1% per Ma and a generation time of 9.2 years, can be translated into a female Ne (Nef) of 43478. However, LAMARC estimates of θ in the varying population size model may also be slightly biased upward, due to correlation with *g*, when only a few loci are analysed (Kuhner et al. 1998). In the analyses with the constant population size model, the highest likelihood estimate of θ per site was 0.004 (95% HPD interval: [0.002, 0.006]; ESS > 850 in all individual replicate runs), which corresponds to a θ of 2.6 for the 648 bp fragment and, assuming a nucleotide substitution rate of 1% per Ma and a generation time of 9.2 years, can be translated into a N_e_f of 21739. This θ estimate was very similar to that obtained using Watterson’s (1975) estimator based on the number of segregating sites: 2.7.

In GENETREE, the analyses using the constant population size model also yielded a ML θ estimate of 2.7 (thus, a N_e_f of 22645), while in the exponential growth model the ML θ estimate was 3.1 (i.e. N_e_f = 26000); the latter model suggested a slow exponential growth rate. However, a LRT did not detect a significant difference in the fit to the data between the simpler constant population size model and the exponential growth model (P = 0.460).

### Time to most recent common ancestor (TMRCA)

Thomson’s method estimates for the TMRCA of the observed genetic variation in *H. hyaena* were 419 ± 112 ka (mean ± standard error; 95% CI: 225-672 ka) based on the 648 bp dataset and a nucleotide substitution rate of 1% per Ma, and 410 ± 160 ka (mean ± standard error; 95% CI: 153-793 ka) based on the 340 bp dataset and a nucleotide substitution rate of 0.93% per Ma (Table 3). Adding the uncertainty in the fossil record and in the *d_xy_* estimate for the divergence between *Hyaena* and *Parahyaena* into the nucleotide substitution rate estimate (i.e. 95% CIs of 0.63-1.45% and 0.51-1.49% for the 648 bp and 340 bp datasets, respectively) yields the following respective 95% CIs for the TMRCA: [155, 1066] and [95, 1447]. In GENETREE, analyses of the 648 bp dataset with the constant population size model and the ML estimate of θ of 2.7 gave a TMRCA estimate, in coalescent time units, of 1.6 ± 0.5 (mean ± standard deviation), which, using a nucleotide substitution rate of 1% per Ma and a generation time of 9.2 years, corresponds to 333 ± 104 ka; approximating the 95% CI by ± 2 standard deviations (Harding et al. 1997) gives 125-542 ka (86-860 ka including uncertainty in the fossil record and in the *d_xy_* estimate for the divergence between *Hyaena* and *Parahyaena* in the nucleotide substitution rate estimation). With the exponential growth model and the respective ML estimate of θ of 3.1, the TMRCA estimate in coalescent time units was 1.27 ± 0.35 (mean ± standard deviation), i.e. 304 ± 84 ka (95% CI: 136-471 ka) (94-748 ka incorporating the aforementioned uncertainty in the nucleotide substitution rate estimate). Analyses of the 340 bp dataset with the constant population size model and a ML estimate of θ of 0.9 yielded a TMRCA estimate in coalescent time units of 1.9 ± 0.8 (mean ± standard deviation), which, using a nucleotide substitution rate of 0.93% per Ma and a generation time of 9.2 years, translates into 279 ± 118 ka (95% CI: 44-515 ka) (28-939 ka considering the uncertainty in the nucleotide substitution rate estimate). Analyses of the same dataset with the exponential growth model and a ML estimate of θ of 1.1 resulted in a TMRCA estimate in coalescent time units of 1.38 ± 0.45 (mean ± standard deviation), i.e. 240 ± 78 ka (95% CI: 83-397 ka) (52-723 ka considering the uncertainty in the nucleotide substitution rate estimate) (Table 3). The fact that GENETREE TMRCA estimates using two very different demographic models were quite similar suggests a minor effect of the underlying demographic model on the estimates. TMRCAs estimates from the BEAST analyses for different combinations of dataset, calibrations, and clock model, are given in Table 3; ESS values were > 3000 for all parameters in all individual runs under the strict clock, and > 200 under a random local clock. Across the different analyses, the TMRCA point estimate for the observed mtDNA variation in striped hyenas ranged from 330-480 ka, with the clock rate point estimate between 1.1% and 1.6% per Ma, hovering around 1.3% per Ma (Table 3), which is quite close to the rate estimates of ≈ 1% that we inferred based on the *d_xy_* distance estimate between *Hyaena* and *Parahyaena*.

**Table 3.**
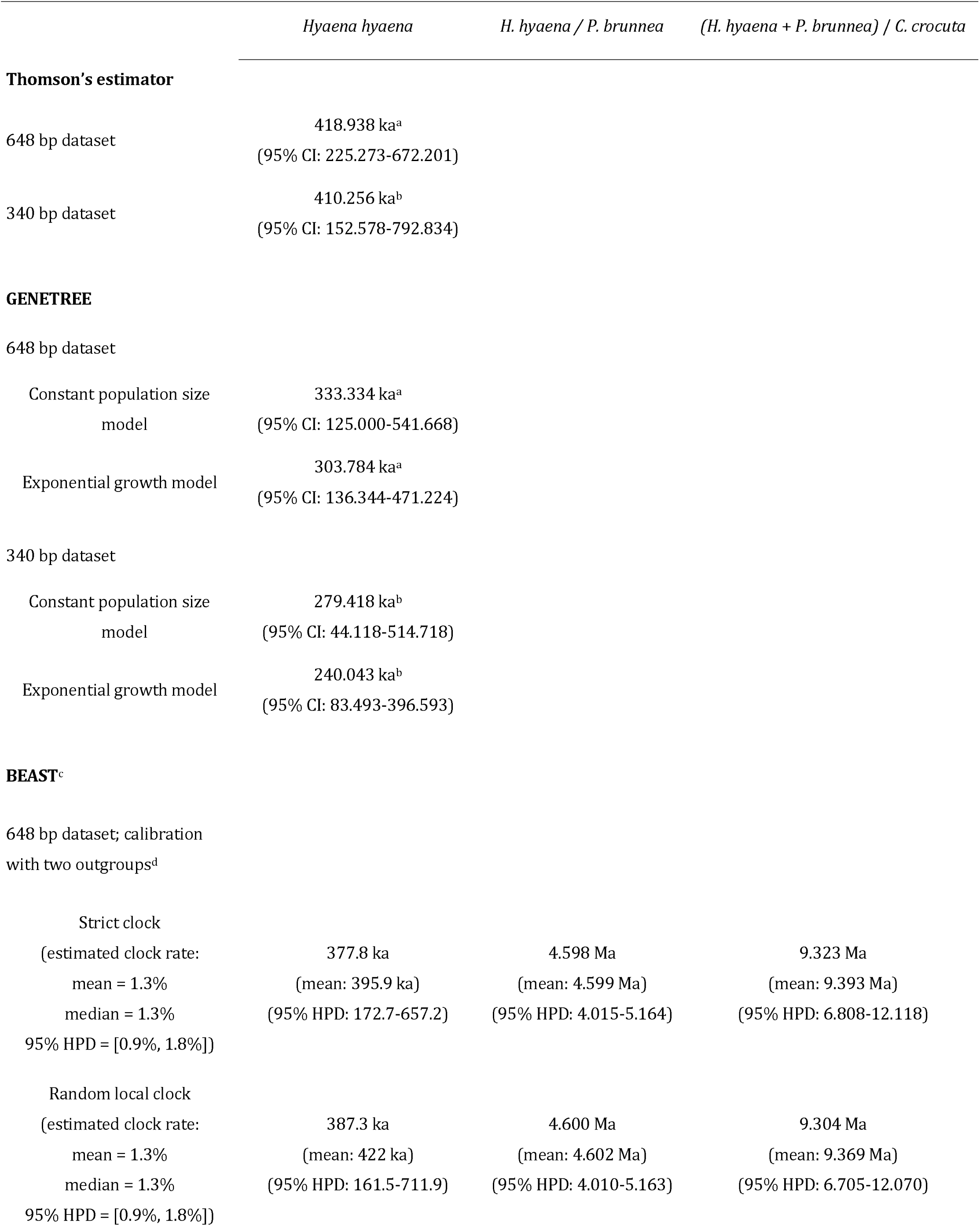
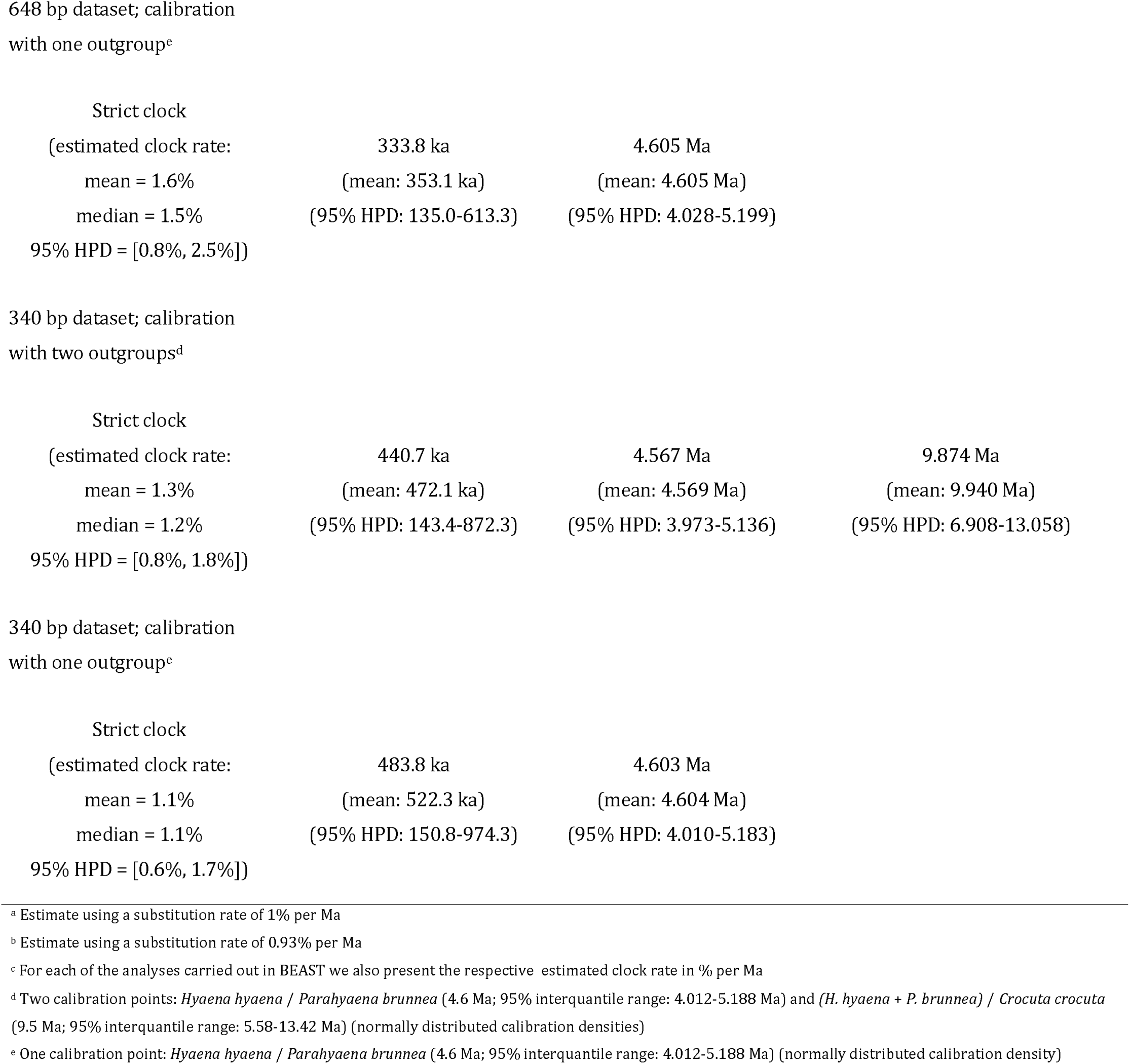
TMRCA estimates for the striped hyena and between striped hyena and the outgroups used for time calibrations in the phylogenetic dating analyses.

|  | *Hyaena hyaena* | *H. hyaena* / *P. brunnea* | *(H. hyaena + P. brunnea)* / *C. crocuta* |
| --- | --- | --- | --- |
| **Thomson's estimator** |  |  |  |
| 648 bp dataset | 418.938 ka<sup>a</sup><br>(95% CI: 225.273-672.201) |  |  |
| 340 bp dataset | 410.256 ka<sup>b</sup><br>(95% CI: 152.578-792.834) |  |  |
| **GENETREE** |  |  |  |
| 648 bp dataset |  |  |  |
| Constant population size model | 333.334 ka<sup>a</sup><br>(95% CI: 125.000-541.668) |  |  |
| Exponential growth model | 303.784 ka<sup>a</sup><br>(95% CI: 136.344-471.224) |  |  |
| 340 bp dataset |  |  |  |
| Constant population size model | 279.418 ka<sup>b</sup><br>(95% CI: 44.118-514.718) |  |  |
| Exponential growth model | 240.043 ka<sup>b</sup><br>(95% CI: 83.493-396.593) |  |  |
| **BEAST<sup>c</sup>** |  |  |  |
| 648 bp dataset; calibration with two outgroups<sup>d</sup> |  |  |  |
| Strict clock<br>(estimated clock rate:<br>mean = 1.3%<br>median = 1.3%<br>95% HPD = [0.9%, 1.8%]) | 377.8 ka<br>(mean: 395.9 ka)<br>(95% HPD: 172.7-657.2) | 4.598 Ma<br>(mean: 4.599 Ma)<br>(95% HPD: 4.015-5.164) | 9.323 Ma<br>(mean: 9.393 Ma)<br>(95% HPD: 6.808-12.118) |
| Random local clock<br>(estimated clock rate:<br>mean = 1.3%<br>median = 1.3%<br>95% HPD = [0.9%, 1.8%]) | 387.3 ka<br>(mean: 422 ka)<br>(95% HPD: 161.5-711.9) | 4.600 Ma<br>(mean: 4.602 Ma)<br>(95% HPD: 4.010-5.163) | 9.304 Ma<br>(mean: 9.369 Ma)<br>(95% HPD: 6.705-12.070) |
| 648 bp dataset; calibration<br>with one outgroup<sup>e</sup> |  |  |  |
| Strict clock |  |  |  |
| (estimated clock rate: | 333.8 ka | 4.605 Ma |  |
| mean = 1.6% | (mean: 353.1 ka) | (mean: 4.605 Ma) |  |
| median = 1.5% | (95% HPD: 135.0-613.3) | (95% HPD: 4.028-5.199) |  |
| 95% HPD = [0.8%, 2.5%]) |  |  |  |
| 340 bp dataset; calibration<br>with two outgroups<sup>d</sup> |  |  |  |
| Strict clock |  |  |  |
| (estimated clock rate: | 440.7 ka | 4.567 Ma | 9.874 Ma |
| mean = 1.3% | (mean: 472.1 ka) | (mean: 4.569 Ma) | (mean: 9.940 Ma) |
| median = 1.2% | (95% HPD: 143.4-872.3) | (95% HPD: 3.973-5.136) | (95% HPD: 6.908-13.058) |
| 95% HPD = [0.8%, 1.8%]) |  |  |  |
| 340 bp dataset; calibration<br>with one outgroup<sup>e</sup> |  |  |  |
| Strict clock |  |  |  |
| (estimated clock rate: | 483.8 ka | 4.603 Ma |  |
| mean = 1.1% | (mean: 522.3 ka) | (mean: 4.604 Ma) |  |
| median = 1.1% | (95% HPD: 150.8-974.3) | (95% HPD: 4.010-5.183) |  |
| 95% HPD = [0.6%, 1.7%]) |  |  |  |

<sup>a</sup> Estimate using a substitution rate of 1% per Ma
<sup>b</sup> Estimate using a substitution rate of 0.93% per Ma
<sup>c</sup> For each of the analyses carried out in BEAST we also present the respective estimated clock rate in % per Ma
<sup>d</sup> Two calibration points: *Hyaena hyaena* / *Parahyaena brunnea* (4.6 Ma; 95% interquantile range: 4.012-5.188 Ma) and *(H. hyaena + P. brunnea)* / *Crocuta crocuta* (9.5 Ma; 95% interquantile range: 5.58-13.42 Ma) (normally distributed calibration densities)
<sup>e</sup> One calibration point: *Hyaena hyaena* / *Parahyaena brunnea* (4.6 Ma; 95% interquantile range: 4.012-5.188 Ma) (normally distributed calibration density)

## Discussion

Topics such as genetic diversity and structure, phylogeography, and evolutionary history of the striped hyena remain poorly investigated (Wilkinson et al. 2024). To date, our knowledge of the mitochondrial genetic diversity and phylogeographic structure of the striped hyena comes essentially from the analysis of a 340-bp *Cyt b* fragment in 13 individuals from across the historical and present distribution of the species, which was part of a ground-breaking study on the evolutionary history of extant hyenas (Rohland et al. 2005). That analysis found low genetic diversity and no evident geographic structure of this diversity. In our study we wanted to investigate the mitochondrial genetic variation of the striped hyena in Algeria, using for this purpose a *Cyt b* fragment encompassing the one analysed by Rohland et al. (2005). It was also our objective, with the addition of samples from the Arabian Peninsula and Iran (regions not covered in Rohland et al. (2005) and where classic morphological subspecies have been described) and using a diverse set of demographic history and coalescence time inference analyses, to conduct a more exhaustive and better-sampled global analysis of the species than in Rohland et al. (2005). The finding that a sample of 24 Algerian striped hyenas was monomorphic for a 753- bp *Cyt b* fragment was striking, and agrees with the notion of low mitochondrial diversity in the species. The apparent absence of phylogeographic structure in the species seemingly extends also to southern Arabia and Iran (Figs. 2 and S2- S4), with for example haplotype sharing between Oman and Iran for the 648-bp fragment (Fig. 2a), which therefore also calls into question the validity of the *sultana* and *hyaena* subspecies.

### Demographic history

The observed patterns of moderate haplotype diversity and low nucleotide diversity (Table 2) are compatible with a historical population bottleneck in the species, followed by demographic growth and the appearance/accumulation of new haplotypes (Grant and Bowen 1998). Moreover, the mismatch distributions for both 648 bp and 340 bp datasets showed a significant fit with expected distributions under a sudden expansion model. Nevertheless, it should be taken into account that tests based on the mismatch distribution have been noted as conservative in rejecting the null hypothesis of population expansion (Ramos-Onsins and Rozas 2002; Fahey et al. 2014). On the other hand, none of the mutation-drift equilibrium tests used rejected the hypothesis of constant population size (Table 2). A reduced number of segregating sites, as is the case here, can limit the power of these tests (Ramos-Onsins and Rozas 2002). In particular, the values of the *F*S statistic were clearly negative, a result consistent with expectations in a scenario of population growth, which may suggest a lack of power in the datasets due to a small number of segregating sites. However, a limited number of segregating sites does not necessarily prevent significant results in those tests (Pilkington et al. 2008; Kvist et al. 2011; Cabanne et al. 2013; Álvarez-Varas et al. 2015; Hassan-Beigi et al. 2022). These tests also have less power if the demographic expansion is historically very recent (e.g. Holocene), especially if it was not very intense (Fu 1997; Ramos-Onsins and Rozas 2002). Rohland et al. (2005) also did not detect a statistical signal of demographic expansion in their data (the authors did not report which analyses and tests of demographic history were performed), but the observed phylogeographic pattern was inferred to be the result of a recent and rapid range expansion, probably within the last 100 ka and possibly as recently as the Holocene, from a small refuge population, most likely located in Africa as, according to Werdelin and Solounias (1991), no Late Pleistocene fossils of striped hyenas are know outside of Africa. The results of the single-tree skyline plot analyses and demographic model fit comparisons all supported the population growth hypothesis. However, these single-tree methods have the limitation of not taking into account uncertainty in the tree topology (Strimmer and Pybus 2001; Pybus and Rambaut 2002); we attempted to mitigate this problem by obtaining the tree to be used by Bayesian analysis and summarizing the posterior into a MCCT, the tree that maximizes the product of the posterior clade probabilities. All the most sophisticated demographic inference methods used (MCMC sampling-based skyline plots, IPHULA, LAMARC, GENETREE) could not reject with high confidence (> 95%) the hypothesis of constant population size. MCMC sampling-based skyline methods may allow to correctly identify demographic trajectories based on single-locus data (Gill et al. 2013), but such datasets may also lack the power to reject the hypothesis of constant demography (Heled and Drummond 2008). For example, although the sample size we analysed cannot be considered small (n = 76), it has been noted in the literature that the BSP method may lack the power to correctly detect demographic expansions using single locus mitochondrial data when the sample size is < 100 (Grant 2015). Again, also for those more complex methods it is obviously preferable that the number of segregating sites is not small (Ho and Shapiro 2011), but they have been applied to datasets with numbers of segregating sites similar to our data, and produced significant results (e.g. Griffiths and Tavaré 1994b, 1999; Kvist et al. 2011; Cabanne et al. 2013; Horreo et al. 2013; Kuo et al. 2014; Álvarez-Varas et al. 2015; Ely et al. 2017). A small number of segregating sites may not be the only challenge for the methods implemented in IPHULA, LAMARC, and GENETREE. Another factor is the assumed population growth model, which in the case of these three methods is the exponential model. It is conceivable that the hypothesis of constant population size may not be rejected with confidence if an exponential growth model is very far from the true demographic history, which could be, for example, one of slow, more or less linear growth, or a complex history alternating recurrent episodes of expansion, stability, and/or decline (e.g. Excoffier and Schneider 1999; Strimmer and Pybus 2001). For example, in the BEAST analysis with the exponential growth coalescent model as tree prior, the marginal posterior distribution of the exponential growth rate parameter included zero, implying that a contrasting model of constant population size could not be rejected. Overall, the observed patterns of haplotype and nucleotide diversity, the mismatch distributions, and the single-tree skyline plot analyses and demographic model fit comparisons supported the hypothesis of population growth, confirming the inference by Rohland et al. (2005). This hypothesis was not statistically significant in the more sophisticated and exacting methods used, probably largely as a result of the limited power provided by a small number of segregating sites.

It was surprising to observe only one haplotype in 24 individuals from Algeria, whereas for example 41 animals from Iran showed eight haplotypes (648 bp fragment; Table 1, Fig. 2a). It is therefore possible that the post-bottleneck demographic growth within Africa was not as rapid and intense as that associated with the colonization of Eurasia. If this was the case, then signatures of demographic expansion will perhaps be easier to detect using large Asian population samples. In the same way that in the human species, signatures of demographic expansion are easier to detect in non-African populations than in African ones (Ingman et al. 2000; Atkinson et al. 2008) or in pooled or global datasets (e.g. Atkinson et al. 2009). Given the typical substitution rates in mitochondrial DNA, it is expected that recent demographic expansions (e.g. postglacial), particularly in organisms whose generation time is not short, may be difficult to detect with data from relatively short mtDNA fragments (e.g. < 1000 bp), even using efficient methods in extracting the information present in the data (Felsenstein 1992). Data from mitogenomes can be very useful in such cases (Keis et al. 2013; Jensen et al. 2018). Future mitogenomic studies will likely help to clarify open questions regarding the nature and age of the Late Quaternary striped hyena demographic and geographic expansion(s) suggested by the currently available mitochondrial data.

### Age of the extant mitochondrial DNA variation

For the estimation of the TMRCA of the sampled genetic variation we also used very different methods, always based on the same rationale that if different analytical approaches yield congruent results, this increases our confidence in them. The Thomson’s estimator is simple and does not require the assumption of a model for the demographic history of the population. GENETREE computes TMRCA likelihood distributions conditional on the gene tree mutation structure, a given value of θ, and the assumed model of population size history. For both approaches the TMRCA estimates were translated into absolute time using the estimated substitution rates for each fragment (340 bp and 648 bp). In contrast, in BEAST the time information to calculate the TMRCA in years was derived from fossil calibrations for the divergence between the striped hyena lineage and the brown hyena and spotted hyena lineages. The TMRCA point estimates of the observed genetic variation ranged between about 300 and 420 ka based on the 648 bp dataset, and between about 240 and 480 ka based on the 340 bp dataset (Table 3); hence there was relative agreement between the estimates from the two datasets. The variability of the estimates was lower for the dataset with longer sequence length and more segregating sites, as would perhaps be expected. For both datasets, estimates were higher with the Thomson’s method (648 bp: 419 ka; 340 bp: 410 ka) and with BEAST (648 bp: 334-387 ka; 340 bp: 441-484 ka) than with GENETREE (648 bp: 304-333 ka; 340 bp: 240-279 ka). This pattern is in line with arguments that GENETREE results should be interpreted as estimated lower bounds for the true values (Hammer et al. 1998; Karafet et al. 1999). In fact, most estimates from Thomson’s method and BEAST tended to hover around 400 ka, coinciding with one of the longest and warmest interglacials of the last 800,000 years (within Marine Isotope Stage 11, MIS 11: 374-424 Ka, Lisiecki and Raymo 2005), with environmental conditions similar to the Holocene (Howard 1997; Kleinen et al. 2014; Tzedakis et al. 2022). Rohland et al. (2005), using the phylogenetic program r8s (Sanderson 2003) and a point estimate of 10 Ma for the divergence between spotted and striped/brown hyenas as a calibration date, estimated the TMRCA of striped hyenas at about 340 ka, which is quite congruent with our estimates. One of our goals in this study was precisely to provide additional estimates, using different methods, for comparison with that one obtained with r8s. For instance, it has been suggested that BEAST may be more accurate than r8s for clade age estimation, particularly with single locus mitochondrial data (Mulcahy et al. 2012).

### Nuclear DNA perspective

A recent whole nuclear genome sequencing study on the evolutionary histories of extant hyenas (Westbury et al. 2021), in which the striped hyena was represented by a single captive individual of unknown geographic origin, indicated a very low genetic diversity also in the nuclear genome of the species, and estimated a demographic growth in the lineage starting ∼ 500 ka that was eventually followed by a sharp decline since ∼ 100 ka. A priority in the near future should be a genome-wide study with a broad geographic sampling throughout Africa and Asia to allow a better characterization of genetic diversity, and the impact on it of the anthropogenic reduction of the species in the last centuries, and a more powerful and informative assessment of population structure and demographic history, as already attempted for its closest extant species, the brown hyena (Westbury et al. 2018).

## Acknowledgments

Carlos Rodríguez Fernandes thanks the support of CE3C and CHANGE through a FCT-Tenure auxiliary researcher contract (FCiência.ID grant 1441) and an auxiliary researcher contract (FCiência.ID grant 0801), and of FPUL for a contract of invited assistant professor. We thank Yassine Righi for a striped hyena sample from Belezma National Park, Algeria. We are extremely grateful to Dr. Helmut Simon for all his advice on TMRCA estimation, and for the NumPy code to compute Thomson’s estimator. We thank Professor Robert Griffiths for his advice on running GENETREE and analysing the results. We thank Dr. Emmanuel Paradis for his advice on how to compute the unfolded site frequency spectrum in his R package PEGAS. We thank Dr. Heiko A. Schmidt for his advice in installing the IPHULA software. Lastly, we would like to thank Mammalian Biology’s Editor in Chief, Professor Heiko Georg Rödel, and two anonymous reviewers for helpful comments and suggestions to improve the manuscript.

## Author contributions

LD and MR performed the laboratory work for genetic analysis. LD and CRF analysed the data and led the writing of the manuscript. All other authors read previous versions of the manuscript and approved the final version. LD, HBH, RHA, SMM, ZA, AC, and PV contributed striped hyena biological samples. YHB produced Figure 1.

## Data availability

Data from this study are available upon reasonable request to the corresponding author.

## Supplementary Information

### Methodological details of analyses using MCMC sampling based skyline methods in BEAST

In the BSP analysis we used four groups and performed analyses with both the piecewise-constant and piecewise-linear models, and for the substitution rate parameter we tested normal, exponential, and uniform prior distributions. The lengths of the MCMC chains were 200 million generations, sampled every 20000th after a pre-burn-in of 20 million. In the EBSP analysis we performed runs with both the stepwise and linear models, and for the substitution rate parameter we tested normal and uniform prior distributions. A uniform distribution was used for the population prior mean. The lengths of the MCMC chains were 200 million generations, sampled every 20000th after a pre-burn-in of 20 million. For the Skyride model, we used time-aware smoothing, a normal prior distribution for the substitution rate parameter, and MCMC chains with 200 million generations, sampled every 20000th after a pre-burn-in of 20 million. For Skygrid, we used 75 grid points, a cutoff value of 1 Ma, a normal prior distribution for the substitution rate parameter, and MCMC chains with 200 million generations, sampled every 20000th after a pre-burn-in of 20 million. For each of the four types of MCMC sampling-based skyline plot analyses, we used Tracer 1.7.2 (Rambaut et al. 2018), with a default burn-in of 10% of the chain length, to assess convergence of the chain to the stationary distribution, plot marginal posterior densities, and obtain estimates and ESS of parameters. To draw the skyline plots we also used Tracer, and the Python script popGraphFromCSV.py (obtained from https://github.com/benb/beast-mcmc/tree/master/doc/EBSP/scripts).

---

**Fig. S1.**
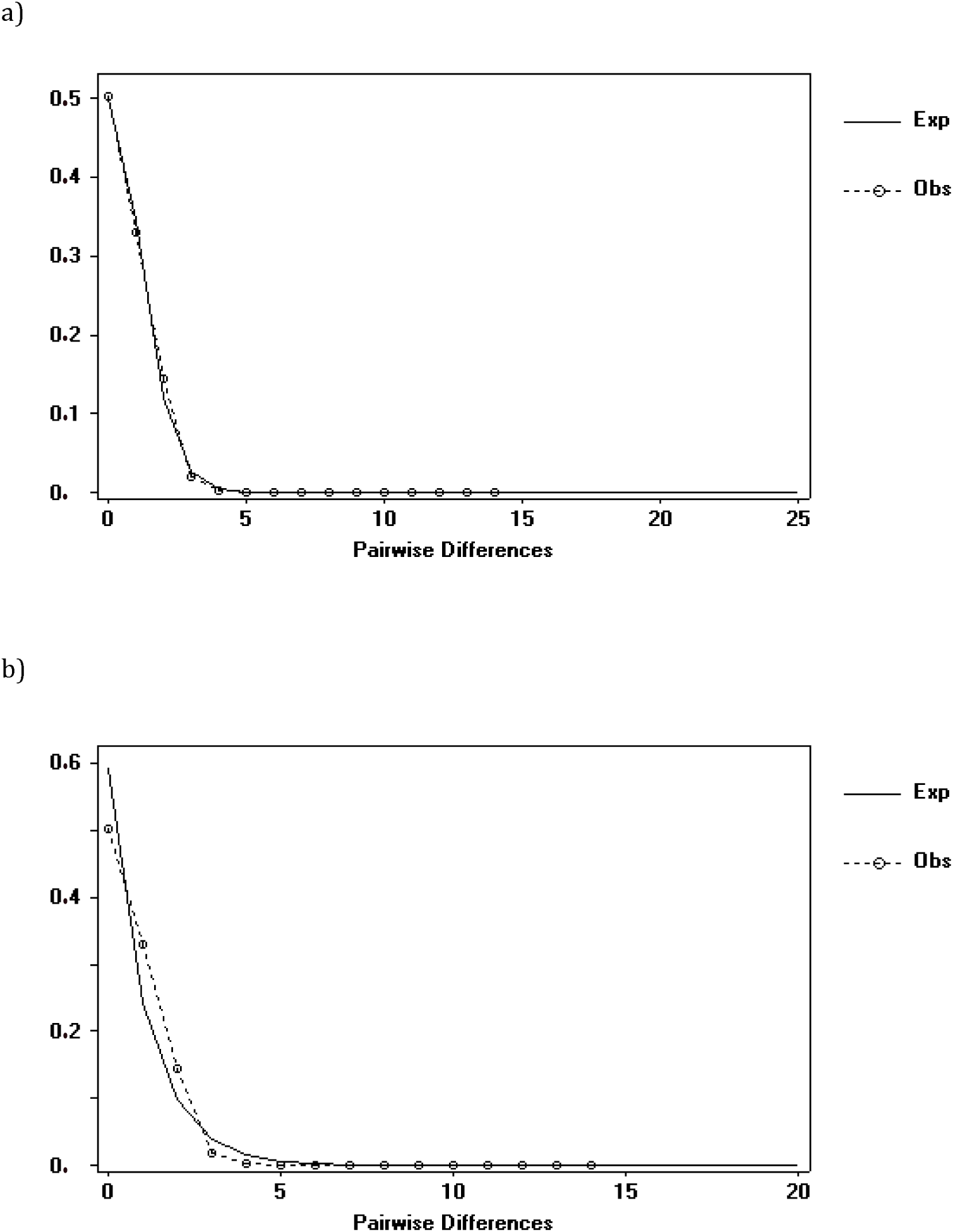
DNASP graphs of mismatch distributions for the striped hyena 340 bp dataset comparing the observed distribution with (a) the expected values under a model of population growth or decline (Rogers and Harpending 1992), and (b) the expected values under a model of constant population size. The x-axis measures the number of pairwise nucleotide differences between sequences, while the y-axis measures the frequency of such pairwise differences in the dataset.

**Fig. S2.**
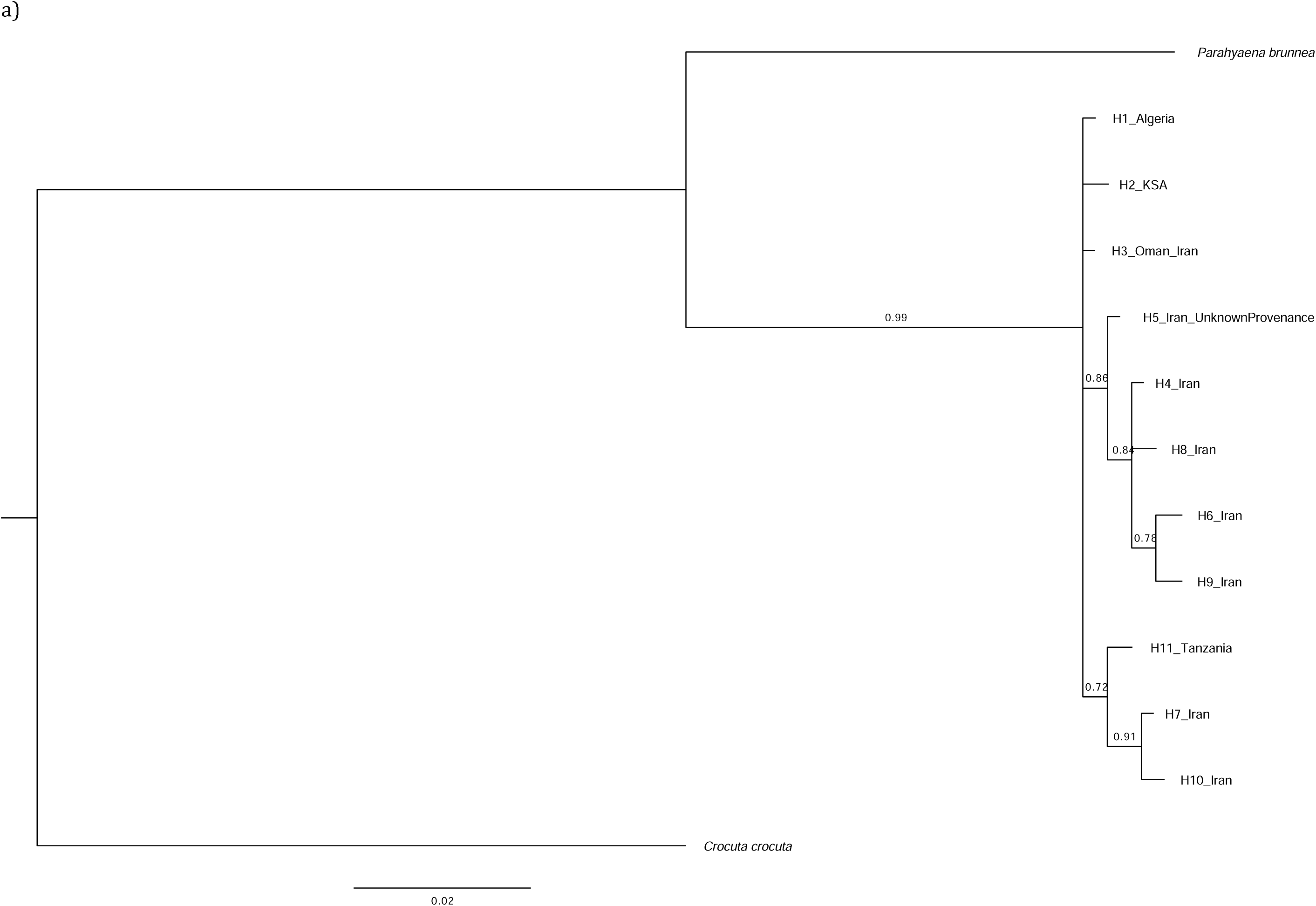
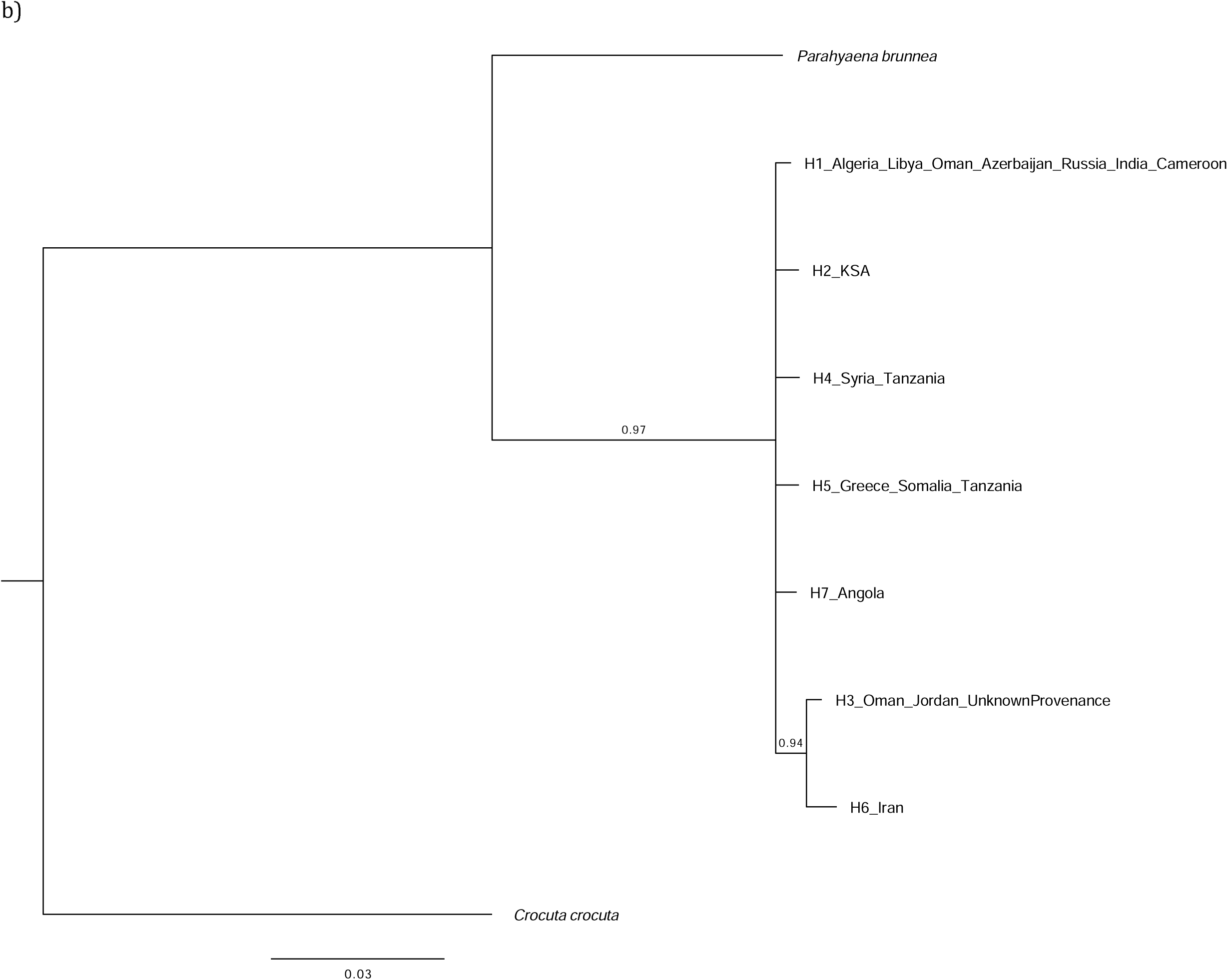
Majority-rule consensus phylograms (cut-off 0.7) from the Bayesian inference analyses of the (a) 648-bp and (b) 340-bp Cyt b datasets, with two outgroups (Parahyaena brunnea and Crocuta crocuta). See Table 1 for haplotype information. Numbers above branches are Bayesian posterior probabilities.

**Fig. S3.**
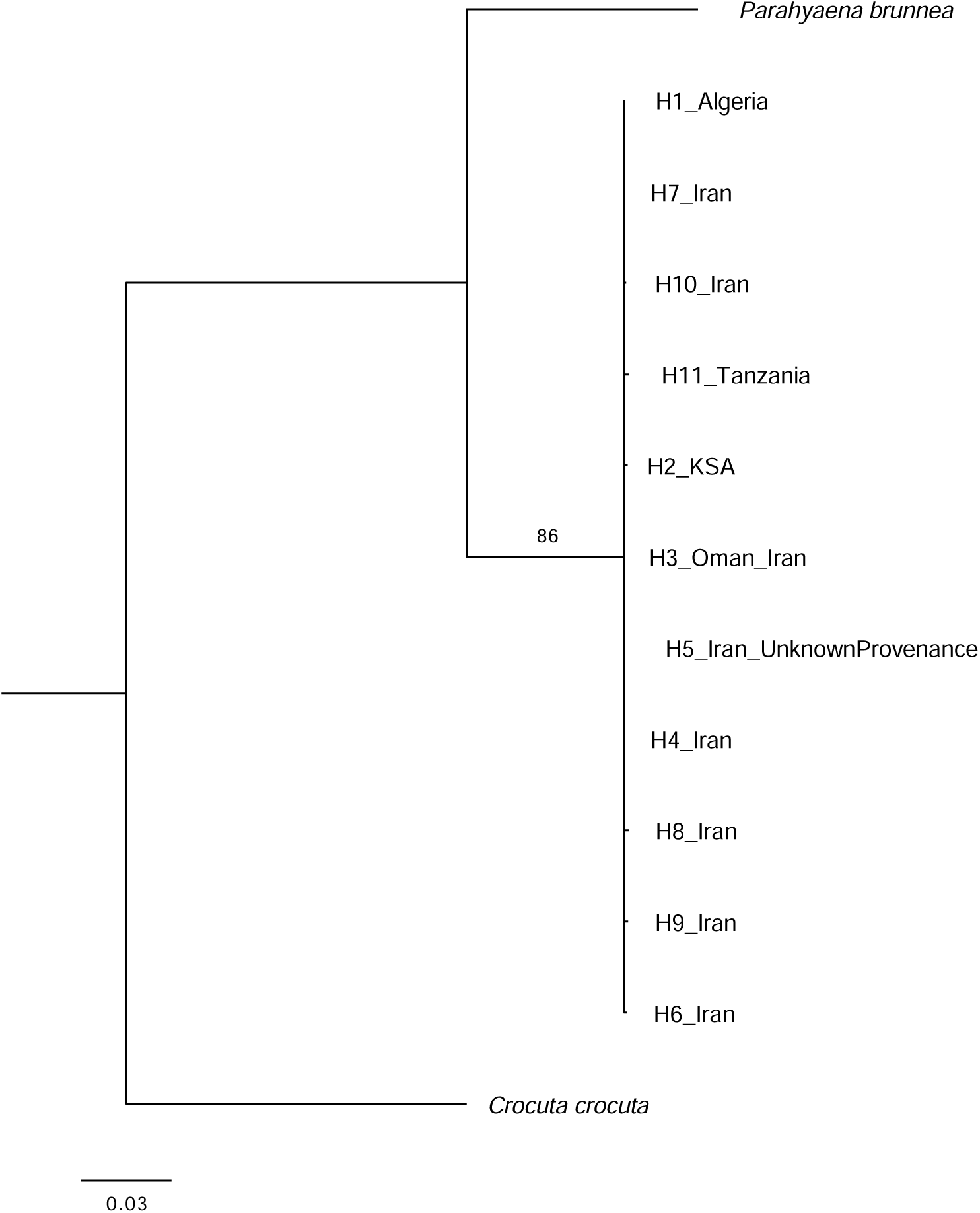
Majority-rule consensus phylogram (cut-off 70%) of the maximum likelihood analysis of the 648 bp dataset with two outgroups (Parahyaena brunnea and Crocuta crocuta). See Table 1 for haplotype information. Numbers above branches are bootstrap values.

**Fig. S4.**
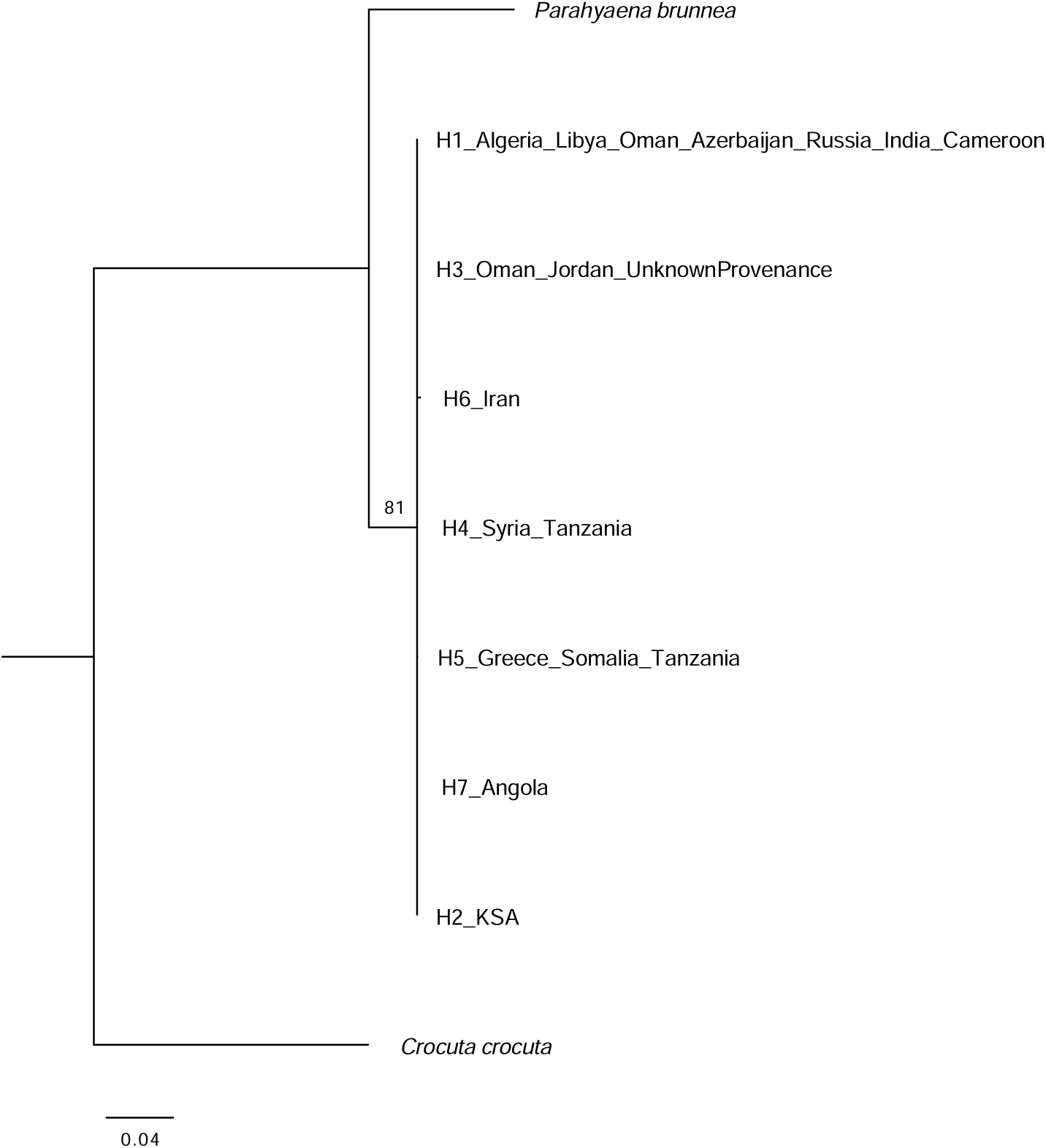
Majority-rule consensus phylogram (cut-off 70%) of the maximum likelihood analysis of the 340 bp dataset with two outgroups (Parahyaena brunnea and Crocuta crocuta). See Table 1 for haplotype information. Numbers above branches are bootstrap values.

**Fig. S5.**
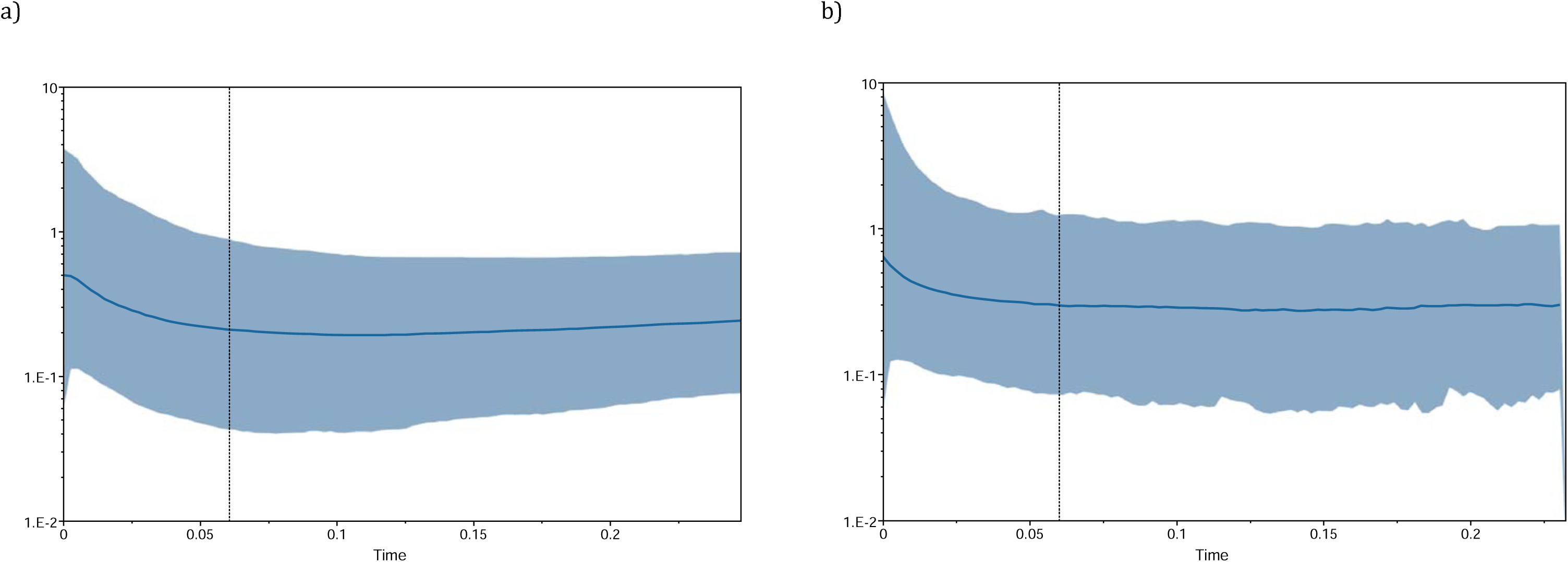
BSPs for the sample of 76 striped hyena 648-bp Cyt b sequences, using the a) piecewise-constant and b) piecewise-linear models, and a normally distributed prior for the substitution rate with mean 0.01 and standard deviation 0.0025 (in substitutions per site per Ma). The thicker solid dark blue line is the median estimate of population size, and the lighter blue shaded region represents the 95% HPD envelope. Maximum time is the root height median, and the black vertical dotted line indicates the lower 95% HPD estimate. The x-axis (in linear scale) measures time in Ma units (going backwards in time from left to right) and the y-axis (in logarithmic scale) measures population size in units of estimated effective population size (N_e_) x generation time in years x 10-6.

**Fig. S6.**
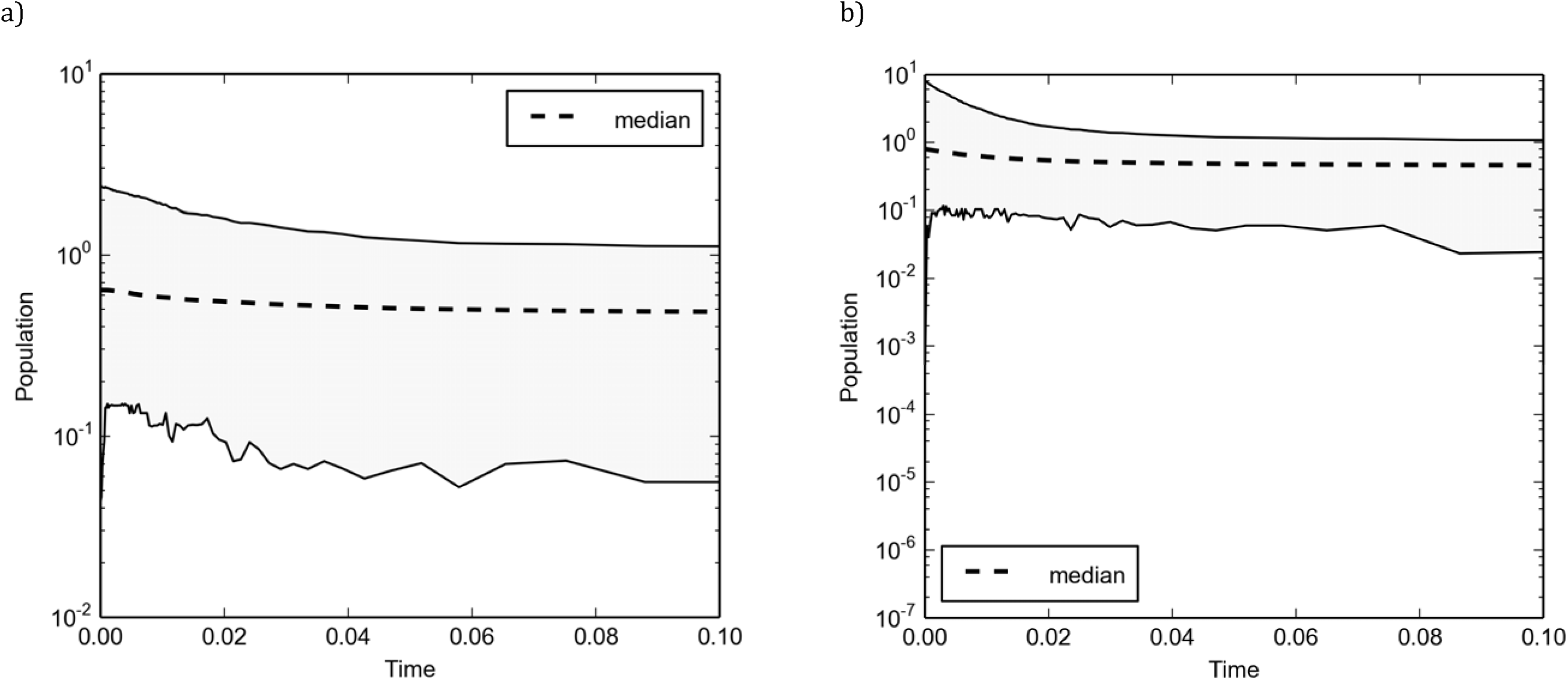
EBSPs for the sample of 76 striped hyena 648-bp Cyt b sequences, using the a) stepwise and b) linear models, and a normally distributed prior for the substitution rate with mean 0.01 and standard deviation 0.0025 (in substitutions per site per Ma). The thicker dashed line is the median estimate of population size, and the shaded region represents the 95% HPD envelope. The x-axis (in linear scale) measures time in Ma units (going backwards in time from left to right) and the y-axis (in logarithmic scale) measures population size in units of estimated effective population size (N_e_) x generation time in years x 10-6..

**Fig. S7.**
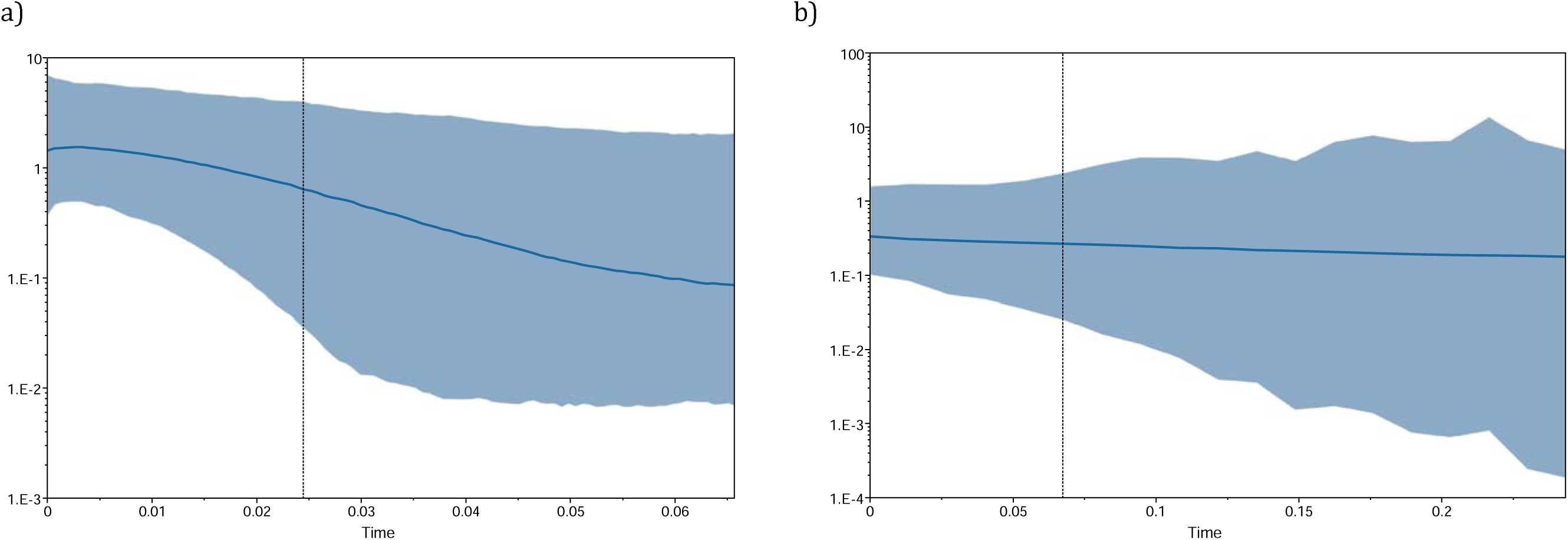
Skyline plots for the sample of 76 striped hyena 648-bp Cyt b sequences, using the a) skyride and b) skygrid methods, and a normally distributed prior for the substitution rate with mean 0.01 and standard deviation 0.0025 (in substitutions per site per Ma). The thicker solid dark blue lines are the median estimates of population size, and the lighter blue shaded regions represent 95% HPD envelopes. Maximum time is the root height median, and the black vertical dotted lines indicate the lower 95% HPD estimates. The x-axis (in linear scale) measures time in Ma units (going backwards in time from left to right) and the y-axis (in logarithmic scale) measures population size in units of estimated effective population size (Ne) x generation time in years x 10^-6^.

**Fig. S8.**
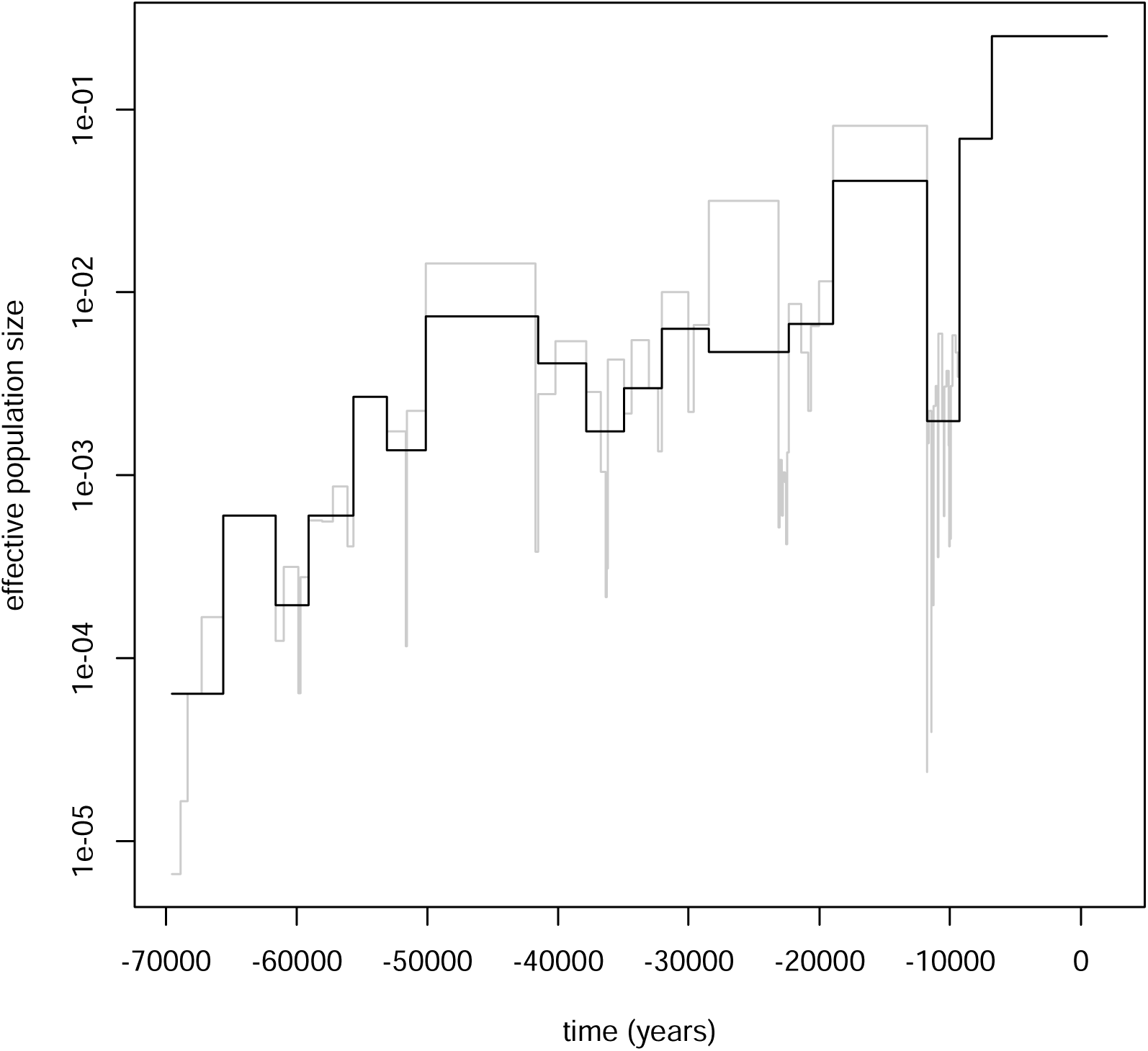
Generalized skyline plot (black line), and classic skyline plot (grey line) in the background. Time is zero at the present.

**Fig. S9.**
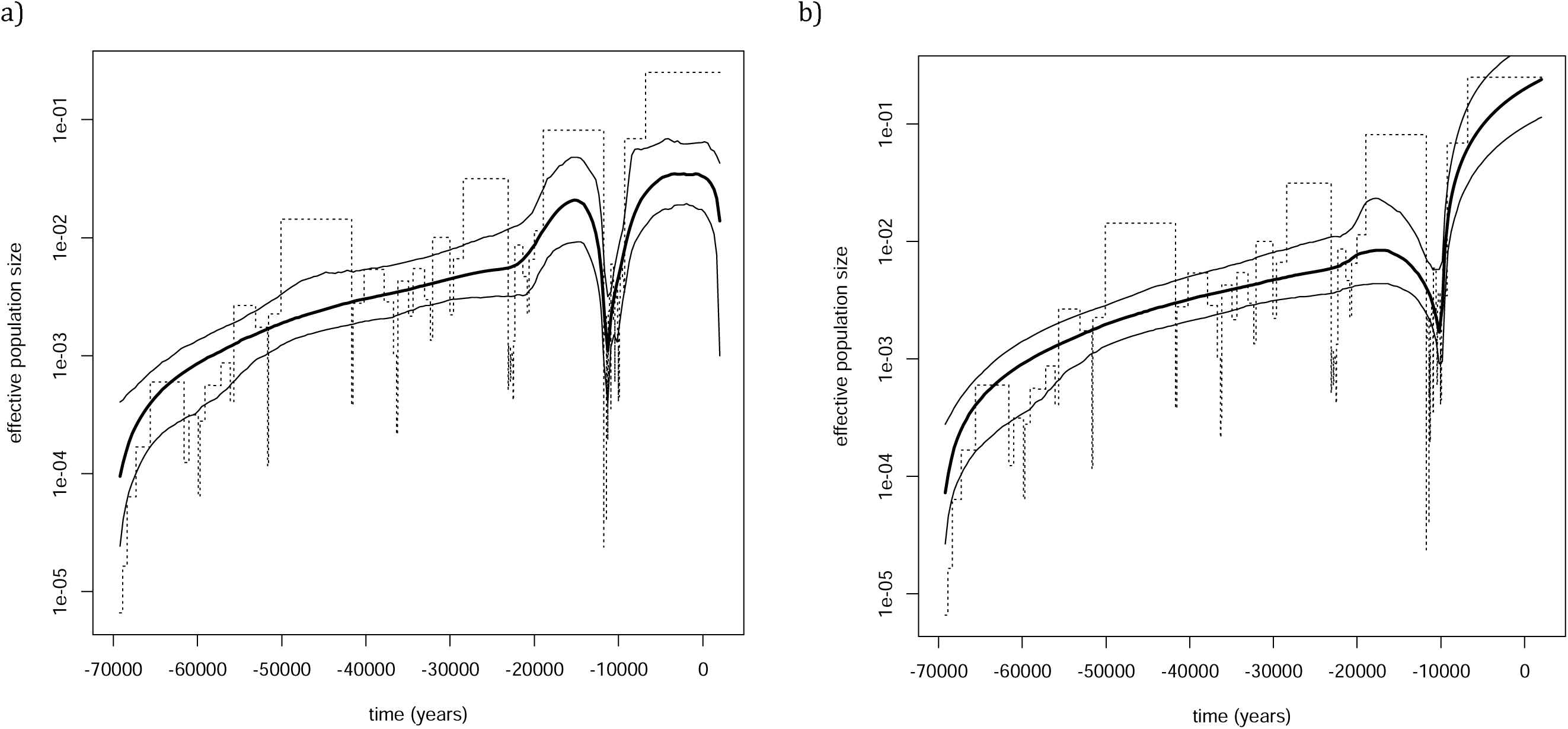
Bayesian MCP plots estimated using a) a constant population size prior and b) a skyline plot prior. The thicker black lines are the median of the posterior distribution of population size, and the thinner lines represent the 95% confidence intervals. In both figures, for comparison, the classic skyline plot is shown in the background (dashed lines). Time is zero at the present.

